# Structural and mechanistic insights into disease-associated endolysosomal exonucleases PLD3 and PLD4

**DOI:** 10.1101/2023.11.20.567917

**Authors:** Meng Yuan, Linghang Peng, Deli Huang, Amanda Gavin, Fangkun Luan, Jenny Tran, Ziqi Feng, Xueyong Zhu, Jeanne Matteson, Ian A. Wilson, David Nemazee

## Abstract

Endolysosomal exonucleases PLD3 and PLD4 (phospholipases D3 and D4) are associated with autoinflammatory and autoimmune diseases. We report structures of these enzymes, and the molecular basis of their catalysis. The structures reveal an intra-chain dimer topology forming a basic active site at the interface. Like other PLD superfamily members, PLD3 and PLD4 carry HxKxxxxD/E motifs and participate in phosphodiester-bond cleavage. The enzymes digest ssDNA and ssRNA in a 5′-to-3′ manner and are blocked by 5′-phosphorylation. We captured structures in apo, intermediate, and product states and revealed a ‘link-and-release’ two-step catalysis. We also unexpectedly demonstrated phosphatase activity via a covalent 3’- phosphistidine intermediate. PLD4 contains an extra hydrophobic clamp that stabilizes substrate and could affect oligonucleotide substrate preference and product release. Biochemical and structural analysis of disease-associated mutants of PLD3/4 demonstrated reduced enzyme activity or thermostability and the possible basis for disease association. Furthermore, these findings provide insight into therapeutic design.

## INTRODUCTION

Nucleic acids are not only the carriers of genetic information but also signal ‘danger’ when mislocalized or presented in aberrant forms. The presence of self or pathogenic DNA or RNA molecules in the cell is detected by various nucleic acid sensors, including Toll-like receptors (TLRs) 3, 7, 8, and 9 in the endolysosomal compartment, and melanoma differentiation-associated protein 5 (MDA5), retinoic acid-inducible gene I (RIG-I), and cyclic GMP-AMP synthase (cGAS) in the cytoplasm [1,2]. All of these nucleic acid sensors trigger nuclear factor kappa-light-chain-enhancer of activated B cells (NF-κB) and type-I interferon pathways to raise a state of alarm in cells, detect self nucleic acids as a signal of cellular damage or stress, or prepare them to resist microbial intruders. However, it is deleterious for cells to be chronically activated, especially when stimulated by nucleic acids from host cells. Host nucleases are therefore important not only in their catabolic functions but also to prevent these sensors from overactivation.

Nucleases cleave phosphodiester bonds between nucleotides. Various endo- and exonucleases have been extensively studied in mammalian cells that have vital functions in regulating innate immunity sensors [3]. For example, DNase II alpha, an acidic endonuclease that hydrolyzes double-stranded DNA (dsDNA) to yield 3′-phosphate and 5′-hydroxyl products, is responsible for degradation of lysosomal DNA from apoptotic cells and processing bacterial genomic DNA for TLR9 activation [4,5]. Three prime repair exonuclease 1 (TREX1) helps prevent autoimmunity by digesting mismatched dsDNA and single-stranded DNA (ssDNA) [6,7]. RNase L, which is induced by interferon, destroys all RNAs upon activation and activates an MDA-5 dependent interferon pathway [8]. In this study, we examined phospholipases D3 and D4 (PLD3 and PLD4), which have recently been shown to be 5′-to-3′ single-stranded DNA/RNA exonucleases localized in endolysosomal compartments [9,10]. Loss-of-function studies have shown that PLD3 and PLD4 are required to prevent inflammation triggered by the host nucleic acid sensors, including TLR9, TLR7, and a sensor coupled to STING (stimulator of interferon genes) [9,10]. Additionally, as PLD3 and PLD4 can cleave substrates as short as dinucleotides, they may also have an important catabolic function.

The structures of nucleases are diverse but their catalytic sites are highly conserved. Catalysis involves nucleophilic attack of the phosphodiester bond in an S_N_2 manner, followed by hydrolysis of the scissile bond. The nucleophiles are usually Ser, Tyr, or His. Penta-covalent intermediates are formed when the nucleic acid substrate is transiently attached to the enzyme. Adjacent acidic or basic residues, and sometimes divalent metal ions (Mg^2+^, Ca^2+^, or Zn^2+^), facilitate stabilization of the negatively charged intermediate [3]. While many endonucleases digest DNA to generate a 5′-phosphate, a few generate a 3′-phosphate group. Based on their enzymatic character, two families of DNase enzymes have been classified [3]: the DNase I family generate products with 5′-phosphate at near neutral pH, many of which require Ca^2+^ or Mg^2+^, whereas the DNase II family generate products with 3′-phosphate in acidic conditions and do not require metal ions.

PLD3 and PLD4 contain 488 and 506 amino acids, respectively, and contain cytoplasmic, transmembrane, and luminal domains. PLD3 and PLD4 belong to the phospholipase D family, as they share two conserved HxK(x)_4_D(E) (abbreviated as ‘HKD/E’) motifs [11–13]. However, the two proteins exhibit a strong exonuclease activity [9,10] instead of the phospholipase activity of PLD1 [14] and PLD2 [15]. PLD3 and PLD4 selectively digest short ssDNA and ssRNA in a 5′ to 3′ manner, and their pH optima are low but differ [9,10], consistent with their endolysosomal location. We confirm here that the pH optimum for PLD4 is considerably lower than that of PLD3.

Some PLD family enzymes (e.g. PLD6, Zucchini, Nuc, and BfiI) exhibit similar biochemical features as PLD3 and PLD4, such as lack of requirement for divalent cations and sensitivity to inhibition by vanadate or tungstate [16]. These enzymes also contain the signature HKD/E motifs. However, the products of these enzymes usually carry a 5′-phosphate and 3′-hydroxyl, whereas PLD3 and PLD4 digestion products have a 5′-hydroxyl and a 3′-phosphate. Structures of PLD family nucleases related to PLD3 and PLD4 have been determined, including endoribonucleases that are essential for primary piRNA biogenesis [17,18], as well as bacterial nucleases Nuc and BfiI, which cleave dsDNA [19,20]. It is then of considerable interest to decipher the enzyme mechanisms of PLD3 and PLD4 with distinct nuclease activities and substrate specificities.

Genome-wide association studies (GWAS) have revealed that *PLD4* is associated with rheumatic diseases such as systemic sclerosis (SSc) [21], systemic lupus erythematosus (SLE) [22], rheumatoid arthritis (RA) [23] and bovine hereditary zinc deficiency (BHZD)-like syndrome [24]. PLD3 is associated with neurodegenerative diseases, such as late-onset Alzheimer’s disease (LOAD) [25–27], spinocerebellar ataxia (SCA) [28], and leukoencephalopathy (LE) [29]. However, the molecular mechanisms of the disease association are not clear.

Here we determined high-resolution crystal structures of mouse PLD3 and human PLD4 in apo, intermediate, and product forms, elucidating the catalytic mechanism of PLD3 and PLD4. Remarkably, we captured an intermediate state in the catalysis where a histidine at the active site was phosphorylated by the 5′-Pi nucleic acid substrate. This observation explains the extremely slow catalysis of 5′-Pi nucleic acids, suggesting a further potential mechanism for the role of PLD3 and PLD4 in innate immunity. Additionally, we observed an extra pair of hydrophobic clamps on PLD4, possibly explaining its slower overall rate of catalysis with nucleic acid substrates. We also demonstrated that disease-associated mutations have a significant impact on the enzymatic activity or stability of PLD3 and PLD4 in vitro and the crystal structures explain the destabilization effect on the enzymes. This comprehensive structural and biochemical study reveals insights in the catalytic mechanism of the PLD3 and PLD4 exonucleases, and may guide structure-based drug design targeting PLD3 and PLD4.

## RESULTS

### Crystal structures of PLD3 and PLD4

PLD3 and PLD4 are type II transmembrane proteins with N-terminal cytosolic tails followed by transmembrane and luminal domains (Figure 1A). Here we expressed and purified the soluble luminal domains of mouse PLD3 (mPLD3) and human PLD4 (hPLD4) (Figure 1A). Human and mouse PLD3 are highly homologous, sharing 94% amino-acid sequence identity (Figure S1). PLD3 is proteolytically cleaved in lysosomes to generate a soluble intraluminal enzyme [30]. To understand the catalytic mechanisms of PLD3 and PLD4, we determined their crystal structures in apo-forms and in complex with a substrate ssDNA, 5′-phosphorylated oligonucleotide (5′-Pi-TTTTT-3′, abbreviated as 5′-Pi-dT_5_) to 2.0–3.0 Å (Table S1). The structures of the two enzymes PLD3 and PLD4 are highly similar, with a Cα RMSD value of 1.1 Å, although their protein sequence identity is only approximately 44% (Figures S1A-1D). Unlike most PLDs, which are homodimers, including Nuc [19], DNase II [31], BfiI [20], and Zucchini [32,33], PLD3 and PLD4 are single-chain molecules composed of two structurally similar domains (A and B) that are related by a pseudo-2-fold symmetry axis (Figure 1). For both enzymes, the two domains are connected by linkers (Figures 1B and 1E). For both PLD3 and PLD4, each domain contains about 200 amino acids that form a seven- or eight-stranded β-sheet flanked by seven helices; domain A contains four antiparallel and three parallel strands, and domain B has five antiparallel and three parallel strands. The two domains form an extensive interface with buried surface areas of 2,143 Å^2^ and 2,076 Å^2^ for PLD3 and PLD4, respectively, which correspond to ∼20% of the total surface area of each molecule. As an intrachain pseudo-dimer, the structures of the two domains are similar with Cα RMSD values of 3.4 Å for PLD3 and 2.4 Å for PLD4 (Figures S1E and S1F). This architecture implies that the enzymes use both domains for catalysis.

**Figure 1.**
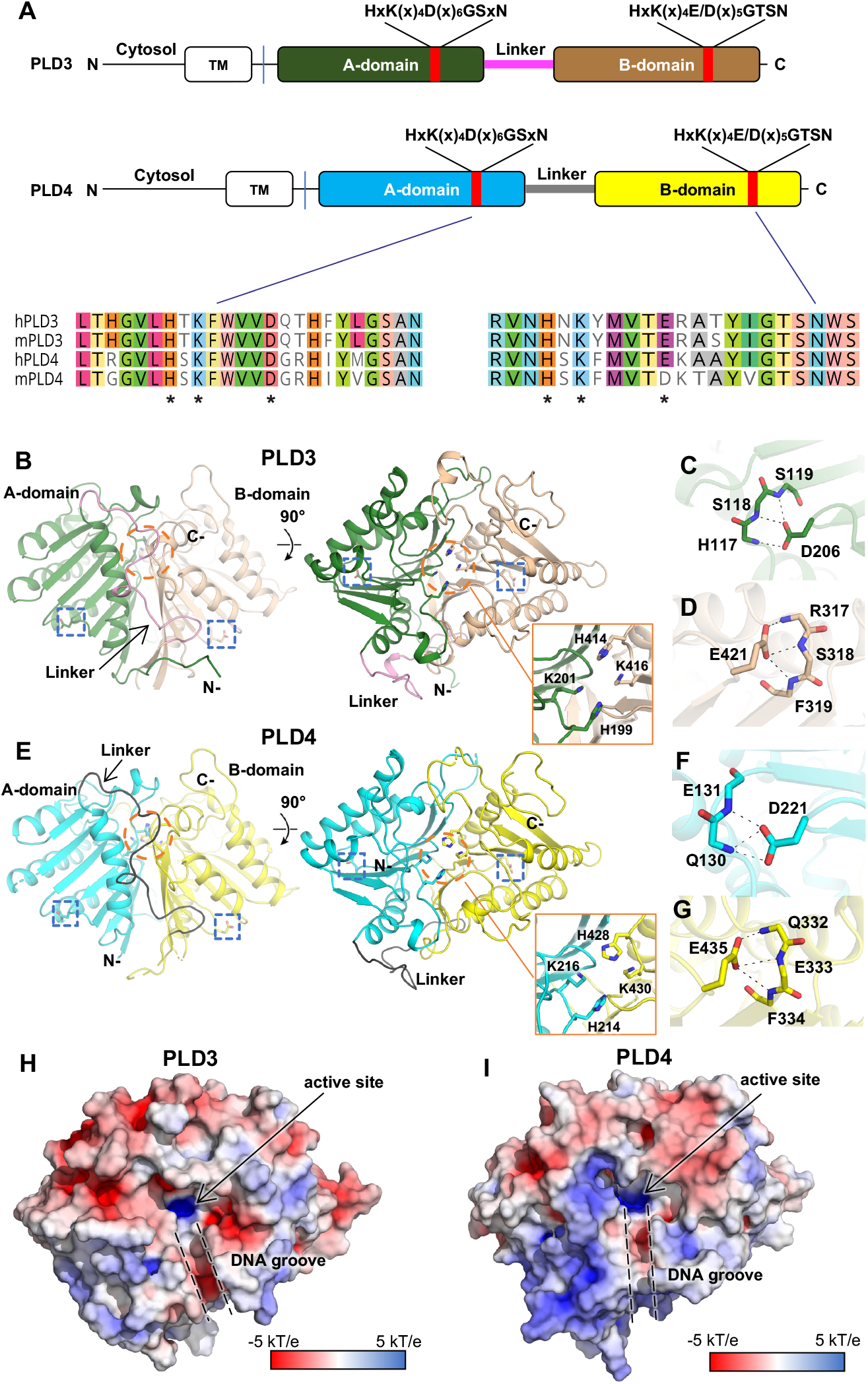
Intrachain-dimeric structures of PLD3 and PLD4. **(A)** Schematic primary structures of PLD3 and PLD4 showing domain structure and HKD motif sequences (marked by asterisks). Prefix “h” represents human and “m” for mouse. TM: transmembrane domain. **(B)** Overall crystal structure of mouse PLD3. The intra-chain pseudodimer is formed by two intra-molecular domains, with A-domain (residues 69–254, green) and B-domain (residues 277–488, brown) linked by residues 255–276 (pink). The active site (orange circle) is formed by both domains. The N- and C-termini are indicated. The blue box highlights HKD motif residues D206 and E421 as zoomed in panels C and D. **(C-D)** Detailed interactions of mPLD3 D206 and E421. The D/E residues form hydrogen bonds with adjacent backbone amides. **(E)** Crystal structure of human PLD4. Similar to mPLD3, the hPLD4 structure contains A (cyan) and B (yellow) domains joined by a flexible linker (black). **(F-G)** Details of interactions of D221, E435 of hPLD4. **(H-I)** Electrostatic potential surface of **(H)** mPLD3 and **(I)** hPLD4. The active sites and the DNA grooves are highlighted with black arrow and dashed lines, respectively. The electrostatic potential surface was calculated with APBS. See also Figure S1 and Table S1.

The HxK(x)_4_D/E consensus sequence exists in both domains of PLD3 and PLD4 (Figures 1A and S1A). The active site is located at the dimer interface, where a pair of histidines and a pair of lysines (Figure 1B and 1E) form a highly basic pocket for accommodating the phosphate group of the nucleic acid substrates, and where a groove formed at the pseudo-dimer interface in PLD3 and PLD4 can accommodate the single-stranded nucleic acid substrates (Figures 1H-1I).

In the HKD/E motifs of PLD3, H199 and K201 in the A-domain oppose their counterparts H414 and K416 in the pseudo-dimer interface, thereby forming the active site of the enzyme (Figure 1B); the acidic residues D206 and E421 likely help stabilize each domain by forming three hydrogen bonds to main-chain amides in spatially adjacent residues (Figures 1C-1D). The HKD/E motifs display similar conformations in PLD4 (Figure 1E), where H214, K216, H428, and K430 form the active site, with D221 and E435 on the sides, potentially stabilizing the protein conformation (Figures 1F-1G). Mutating both D/E to A reduces the protein yield to zero (Figure S2A), supporting the proposed role of the acidic residues in the stability of the PLD proteins, whereas mutation of both H to A had none to less effect on yield. Interestingly, we observed a tartrate molecule in the active site of mPLD3-apo structure (Figures S1G-1H), presumably from the crystallization solution that binds in the anion binding site. This structure with the bound ligand may suggest a starting point for inhibitor design.

### Analysis of 5′-phosphorylated DNA co-crystallized with PLD3 or PLD4

We previously found that mouse PLD3 and PLD4 are specific to 5′-OH nucleic acid substrates [9]. Using gel-based nuclease assays, here we confirmed that mouse PLD3 and human PLD4 exhibited cleavage activity against ssDNA substrates carrying 5′-OH, whereas the enzymes did not appear to cleave ssDNA carrying 5′-phosphate (Figure 2A). hPLD4 with both active-site histidines mutated to alanines lacked catalytic activity, reflecting the important function of these residues (Figure 2A). To generate a co-crystals of PLD3 and PLD4 with their substrates, we crystallized mPLD3 and hPLD4 with a 5′-phosphorylated ssDNA (5′-Pi-TTTTT-3′). The PLD4 co-crystal was grown for 14 days before being tested for x-ray diffraction. The structure illustrated that one histidine and one lysine residue from each HKD/E motif (H214 and K216; H428 and K430) are involved in the interactions with the substrate (Figure 2B). E242 interacts with the histidine of the second HKD/E motif (H428) and helps position it in the active site. Similar to a previously reported PLD family member Nuc, upon substrate binding, the histidine of the second HKD/E motif (H428) appears to function as a nucleophile, while the first histidine (H214) as a general acid protonating the leaving group [19]. The structure clearly illustrates that three nucleotides are bound to the enzyme, where the first base (dT_1_) is clamped between two hydrophobic residues L183 and F423, and the third base (dT_3_) is sandwiched by V212 and F348 (Figure 2B). The structure unexpectedly demonstrated that the second histidine (H428) at the active site of hPLD4 was phosphorylated, suggesting that the 5′-Pi-ssDNA was cleaved between the 5′-phosphate group and the ssDNA by H428, leaving a covalent 3-phosphohistidine (pHis) (Figure 2B), which explains the inhibitory effect of 5′-Pi-ssDNA on the enzyme activity of the PLD. The active-site histidine in the A-domain (H214) was not observed to be phosphorylated.

**Figure 2.**
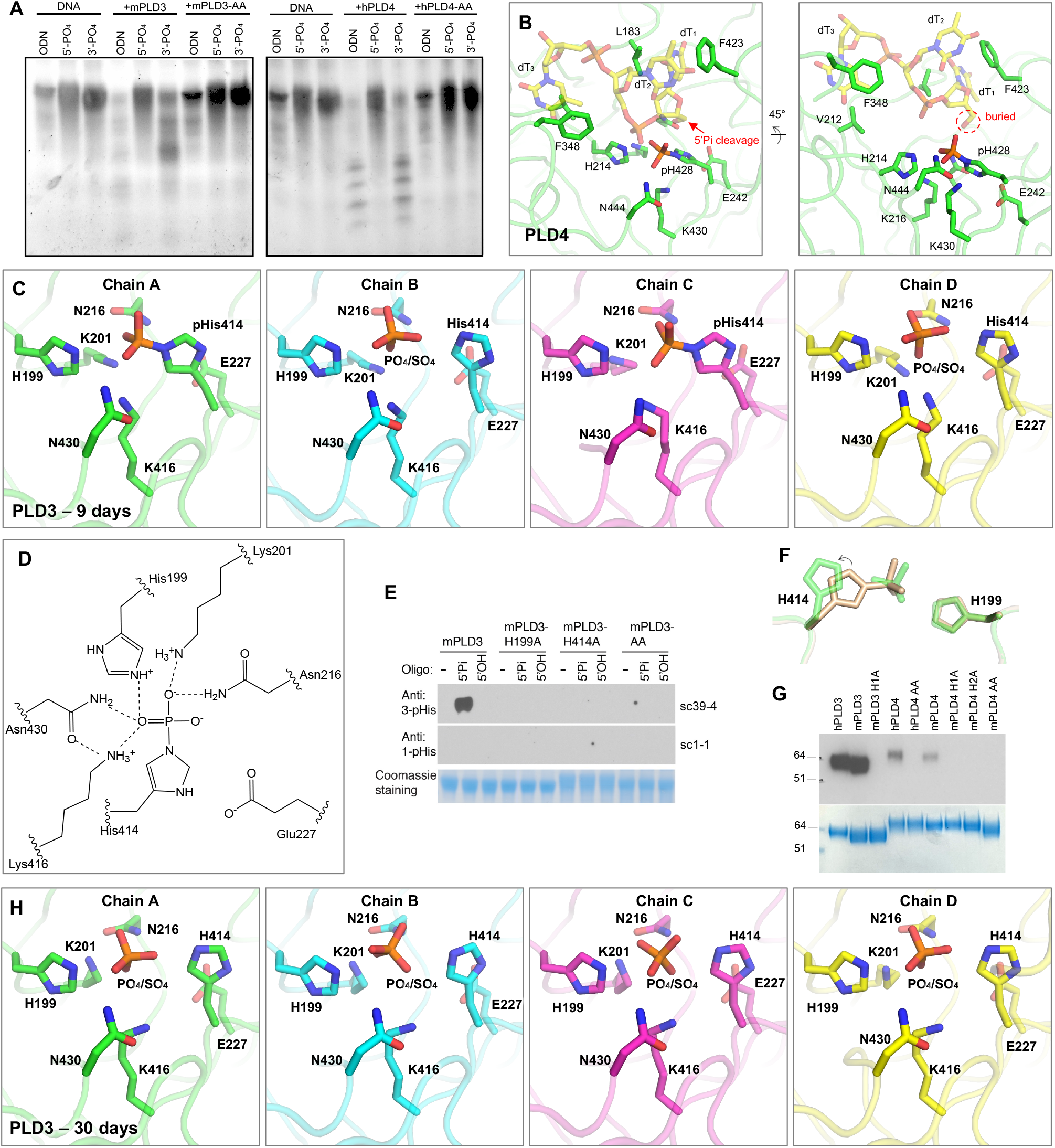
Analysis of co-crystals of PLD3 and PLD4 with phosphorylated ssDNA substrate reveals transfer of 5′-phosphate to the 3-position of an active-site histidine and as well as phosphate release. **(A)** Gel-based assay shows digestion by PLD3 and PLD4 of single-stranded DNAs. ODN: a 55mer oligodeoxynucleotide with no phosphorylation at either 5′ or 3′ end; 5′-PO_4_: a 55mer oligodeoxynucleotide with phosphorylation at only the 5′ end; 3′-PO_4_: a 55mer oligodeoxynucleotide with phosphorylation at only the 3′ end. **(B)** Crystal structure presenting the active site of hPLD4 (green) co-crystallized with 5′-Pi-ssDNA (represented by transparent yellow sticks). A putative phosphodiester cleavage site is indicated by a red arrow. Hydrogen bonds and salt bridges are represented by black dashed lines. **(C)** Crystal structure of mPLD3 co-crystallized with 5′-Pi-ssDNA for nine days. Four copies of molecules were found in each asymmetric unit. Covalent pHis residues were found in chains A and C, and free phosphates or sulfates in chains B and D. **(D)** A hydrogen-bonding network in the active site of mPLD3-pHis as shown in panel C, chain A. The dashed lines represent atoms within 3.4 Å. **(E)** Western blot analysis of histidine phosphorylation of PLD3 incubated with oligonucleotides carrying or lacking a 5′-phosphate. Antibodies specific to 3-pHis (sc39-4) and to 1-pHis (sc1-1) [34] were used. **(F)** Comparison of active site histidines before (sand color) and after phosphate cleavage (green). **(G)** Covalent phosphate is transferred from the 5′-phosphorylated oligonucleotide to PLD3 and PLD4. Briefly, 5′-^32^Pi-ssDNA was incubated with PLD3 or PLD4 in exonuclease buffers and radioactive signals were observed in the proteins after SDS-NuPAGE gel separation. AA/H1A/H2A: both/first/second catalytic histidine(s) mutated to alanine. **(H)** Crystal structure of mPLD3 co-crystallized with 5′-Pi-ssDNA for 30 days. See also Figures S2, S3, and S7.

Here we successfully captured the nucleic acid catalytic product, now carrying a 5′-OH, likely indicating that the hPLD4-pHis blocked the subsequent catalysis. The resulting hydroxyl group of the first nucleotide was adjacent to the phosphorylated hPLD4-H428, indicating the cleavage site of the 5′-Pi-ssDNA (Figure 2B). The 5′-terminus of the ssDNA substrate is completely buried by the enzyme, leaving no room for binding a nucleotide that is extended upstream (Figure 2B) and explaining in part the lack of endonuclease activity [9,10]. These results suggest that PLD4 also has 5′-polynucleotide phosphatase activity in addition to exonuclease activity and show how oligonucleotide substrates fit in the active site.

mPLD3 was also co-crystallized with the same ligand for nine days, and crystals that were formed were tested for diffraction. Like the hPLD4 structure, the mPLD3 also demonstrated a catalytic intermediate state capturing a pHis, where the histidine in the second HKD/E motif (H414) is covalently linked to phosphate (Figures 2C-2D). K201 and K416 are juxtaposed and form salt bridges with the pHis. N216 and N430 are also located on opposite sides of the phosphate and hydrogen bond to the phosphate in the pHis intermediate state. H199 is also opposite the 3-pHis and forms an H-bond/salt bridge with the phosphate (Figures 2C-2D). Interestingly, in the four copies of mPLD3 in the asymmetric unit of the crystal (Figure S3A), H414 was phosphorylated in only two copies (chains A and C) (Figures 2C and S3B). No nucleic acid was observed as PLD3 appears to lack the hydrophobic residues that clamp oligonucleotides in PLD4 (see below). To confirm the phosphorylation observation biochemically, we applied 1-pHis or 3-pHis specific monoclonal antibodies [34] to detect phosphorylation of PLD3 after incubation with 5′-Pi- or 5′- OH-ssDNA substrates (Figure 2E). The results showed that (1) only coincubation with 5′-Pi-ssDNA, but not 5′-OH-ssDNA, resulted in a phosphorylated histidine on PLD3, suggesting that the 5′-Pi of the ssDNA was transferred to the histidine; (2) H199 and H414 are both in the active site, but only the second histidine is phosphorylated; (3) only the 3-pHis but not the 1-pHis monoclonal antibody recognizes the pHis, which corresponds with the observation of 3-pHis in the crystal structures.

The phosphorylated and non-phosphorylated H414 in PLD3 adopt slightly different rotamers, suggesting the histidine side-chain rotates to become covalently attached to the phosphate group when it is located in the basic binding pocket (Figure 2F). Incubation of PLD3 or PLD4 with 5′- ^32^[P]-phosphorylated oligonucleotide followed by SDS-NuPAGE gel electrophoresis revealed radioactive labeling of the enzymes, while no His phosphorylation was observed when one or both active site histidines was mutated, further suggesting the phosphate on the pHis residue was derived from the oligonucleotide substrate (Figure 2G).

Due to the relatively unstable nature of pHis [35], we anticipated that the covalent link may be slowly hydrolyzed. In fact, in the copies where H414 were not phosphorylated (chains B and D), electron density for a tetrahedral anion was found at the active sites that could have been derived from hydrolyzed phosphates as catalytic products, or from sulfates from the crystallization solution (Figures 2C and S3B). Additionally, in contrast to the covalent pHis observed in the mPLD3/5′-Pi-ssDNA crystals harvested after nine days of crystallization, a crystal harvested at 30 days showed that no phosphates were covalently bound to the His (Figures 2H and S3C). The structure of the covalent pHis explains the inhibitory effect of 5′-Pi ssDNA on PLD3/4 enzymatic activity.

### Quantitative enzyme assay confirms 5′-phosphorylated DNA is an inhibitor of PLD3 and PLD4

According to the gel-based assay (Figure 2A), PLD3/4 exhibited minimal enzyme activity against 5′-phosphorylated ssDNA. To quantitatively measure PLD3/4 enzyme activity at molecular and cellular levels, we used a cell-based assay reported before [9], and developed an enzyme-based assay. In the enzyme-based fluorophore-quencher assay, iFr-5-dT, a short thymidyl pentamer with an FAM group on the second thymine and Iowa Black^®^ quencher on the 3′ end, was designed to mimic ssDNA substrate (Figures 3A and S4A). Structural modeling demonstrates that the FAM molecule does not sterically clash with the enzyme (Figure S4B). Cleavage of this fluorogenic substrate by PLD3 and PLD4 could be readily quantified. Optimum substrate and enzyme concentration and reaction conditions were determined (Figure 3B). PLD3 digested substrate much faster and at a lower enzyme concentration than PLD4. Here, our fluorophore-quencher enzyme assay further illustrated that 5-fold excess 5′-phosphorylated ssDNA (5′-Pi-dT_5_) inhibited digestion of iFr-5-dT by PLD3 and PLD4, whereas non-5′-phosphorylated ssDNA (5′-OH dT_5_) at the same concentration did not (Figure 3C). At much higher concentrations (10-250 fold excess), 5′-OH dT_5_ had some inhibitory effect that may in part reflect competition with iFr-5-dT for the basic active site, but this effect was much less pronounced than for 5′-Pi-dT_5_ (Figure S5B). These data show that the nuclease activity of PLD3 and PLD4 can be inhibited by 5′-phosphorylated oligonucleotide.

**Figure 3.**
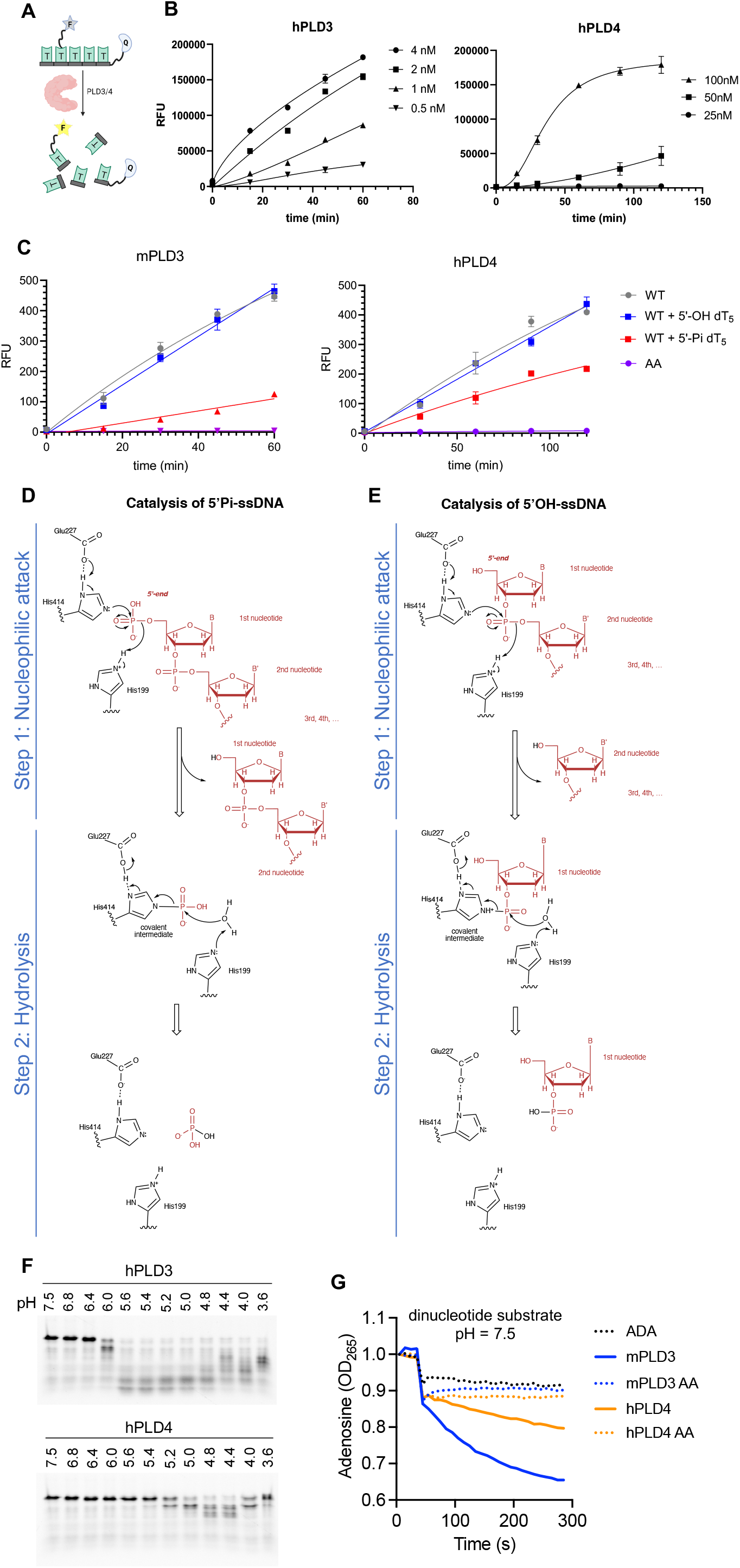
Proposed exonuclease and phosphatase reaction mechanisms and inhibitory effect of 5′ phosphorylation. **(A-B)** Enzyme activity characterization of PLD3 and PLD4. **(A)** Scheme of the fluorophore-quencher assay for PLD3 and PLD4 nuclease activity quantitation. **(B)** Kinetic curves under optimum conditions for PLD3 (MES buffer pH 5.5) and PLD4 (NaAc buffer pH 4.7). Enzyme concentrations were varied as indicated. Substrate concentration was 2 μM for all reactions. **(C)** 5′-Pi-ssDNA inhibits the enzyme activity of PLD3 and PLD4. Analysis of inhibition of PLD3 and PLD4 nuclease activity was conducted using the fluorophore-quencher-labeled ssDNA 5′-OH-ssDNA substrate in the absence (WT) or presence of 5-fold excess amount (10 μM) of unlabeled 5′-Pi-dT_5_ (red) to test its inhibitory effect, while unlabeled 5′-OH-dT_5_ (blue) was used as a control. AA: PLD3/4 with both catalytic histidines mutated to alanines. **(D-E)** Proposed catalytic mechanisms of PLD3 for ssDNA substrates carrying **(D)** 5′-phosphate and **(E)** 5′-hydroxyl groups. Components from the substrates are shown in red. The numbering of mPLD3 residues was used to represent the active-site residues. The first step is nucleophilic attack, where H414 attacks the phosphodiester bond to form a covalent phospho-histidine intermediate (pHis) with the help of E227. The second step is hydrolysis, where H199 is involved in hydrolysis of the pHis intermediate via activation of a water molecule. An analogous mechanism is proposed for PLD4. **(F)** Exonuclease activity of human PLD3 and PLD4 at different pH values. Analysis of digestion of ssDNA 55SUB containing a 3′-FAM group by either human PLD3 or PLD4 at the indicated pH at 37°C for 2 hours at a molar ratio 1:100 enzyme:substrate. **(G)** A dinucleotide assay (5′-OH UpA substrate) at pH 7.5. This assay is coupled with adenosine deaminase which only works on the adenosine when released. ADA alone is shown on the graph where no PLD is added (shown as ‘ADA’). The absorbance shift of adenosine to inosine was measured at 265 nm. AA: a variant of PLD3/4 with both active-site histidines mutated to alanines. See also Figures S4 and S5.

In the cell-based assay, HEK293Blue^TM^ hTLR9 reporter cells with PLD3 knockout (KO) were used to report NF-κB stimulation by CpG-containing oligodeoxynucleotides (ODN), 2006PD (phosphodiester bond) or 2006PS (phosphorothioate bond). Within this assay, reconstitution of PLD3 or PLD4 digests 2006PD and reduces the signal, whereas 2006PS is resistant to PLD3/4 cleavage (Figure S5A) [9]. In contrast to the fluorophore-quencher enzyme assay at the protein level, inhibition by 5′-phosphorylated ssDNA was not observed in the cell-based assay, possibly due to the presence of intrinsic phosphatases (Figure S5C).

### Catalytic mechanism of PLD3 and PLD4

Mutating either or both of the histidines to alanines in the active site completely abolishes enzyme activity, suggesting that both histidines are critical for catalysis (Figure S2B). The structural and biochemical data allow us to propose models for the catalytic mechanisms for the phosphatase and nuclease activities. The two histidines in HKD motifs are essential for the nucleophilic attack and subsequent cleavage of phosphodiester bonds. In the hPLD4 and the tartrate-bound PLD3 structures, a glutamate (E242/E227) H-bonds with the N1 atom of the histidine (H428/H414) in the second domain and may play a role in mechanism of nucleophilic attack by the histidine (Figures 2B and S1G-H) as in other nucleases [19,20,32,33]. Our structures of PLD3 and PLD4 demonstrate nucleophilic attack and subsequent cleavage of the 5′-Pi of the 5′-Pi-ssDNA substrates, where the histidine in the second HKD motif (hPLD4: H428 or mPLD3: H414) attacks the 5′-Pi of the substrate forming a covalent pHis intermediate, followed by a hydrolysis step (Figure 3D).

Based on this observation, we propose a possible catalytic mechanism for the exonuclease activity on the 5′-OH ssDNA substrate (Figure 3E). The first phosphate group of the 5′-OH-ssDNA is located at the 3′ position of the first nucleotide. The second of the two histidines in the active site also makes a nucleophilic attack on the phosphodiester bond between the first and second nucleotides of the substrate nucleic acid to form a histidine-linked covalent intermediate; the other histidine can then act as a general acid protonating the oxygen atom of the leaving group (step 1). A similar covalent bond with the 3′-Pi of the nucleotide substrate as a transient intermediate has been observed in other DNA enzymes, e.g. a 3′-phosphotyrosyl bond is formed as a transient intermediate for the catalysis of topoisomerase [36,37]. Subsequently, the covalent phospho-nitrogen (P-N) intermediate is hydrolyzed by a water molecule with the aid of the histidine in the first HKD motif (hPLD4: H214 or mPLD3: H199), which deprotonates the water to release the 3′- phosphate nucleotide from the enzyme (step 2). In short, the first nucleotide is cleaved off from the nucleic acid substrate and covalently links to a histidine of the enzyme through a P-N bond in the first catalytic step, and is further hydrolyzed by an activated water molecule in the second step. The overall catalysis exhibits a ping-pong mechanism as for PLD1, PLD2, and BfiI, i.e. link-and-release of a nucleotide [19,38,39].

We have previously demonstrated that the endolysosomal enzymes PLD3 and PLD4 exhibited optimal nuclease activity at low pH [9,10]. Here we confirmed that both enzymes require acidic conditions for the overall enzymatic activity, while minimal activity was observed in the assay with a longer nucleic acid substrate that contains 55 nucleotides at neutral pH (Figures 3F and S2C). However, it is not clear which step of the two-step catalysis (Figure 3E) requires the acidic environment. To address this question, under neutral pH, we tested the catalysis of a dinucleotide 5′-OH-UpA using an enzyme assay coupled with adenosine deaminase that only reports signal for single nucleotide adenosines, that is to say, a positive signal will arise only if step 1 in the catalysis is achieved, which releases the second nucleotide (adenosine) (Figure 3E). Interestingly, the experiment showed that the enzymes were able to cleave the dinucleotide even at neutral pH (Figure 3G). This result suggests that the first step of the catalysis (nucleophilic attack), which only cleaves the first nucleotide, can take place at both low and neutral pH. However, the second hydrolysis step can only appear to proceed at low pH to cleave the covalent phospho-intermediate. It remains to be determined why PLD4 requires a lower pH for activity than PLD3 (Figure 3F).

Many PLDs have been reported as metal-ion-independent nucleases [3]. Here we confirm that the enzymatic activity of PLD3 and PLD4 are independent of Ca^2+^ or Mg^2+^. The addition of CaCl_2_/MgCl_2_ or EDTA does not affect enzyme activity, indicating that divalent metal ions (Ca^2+^ or Mg^2+^) are not required for PLD3/4. Interestingly, Fe^2+^, Cu^2+^, and to a lesser extent Zn^2+^, inhibits PLD3/4 enzyme activity at 2 mM, possibly by chelating the catalytic His residues [40] to compete with the substrate (Figure S2D).

### PLD4 has an additional hydrophobic clamp to confer substrate specificity

We also compared the structures of PLD3 and PLD4. A conserved pair of hydrophobic residues F333/348 and V197/212 (mPLD3 and hPLD4 numbering, respectively) clamp the third base of the catalytic product (Figures 4A and S1A). Mutating both residues to alanine abolishes the activity of both PLD3 and PLD4 (Figure 4B). PLD4 has an additional pair of hydrophobic clamp residues L183/F423 that secure the first base of the catalytic product (Figure 4A). In contrast, PLD3 lacks one side of the clamp, where L183 in PLD4 corresponds to G170 in PLD3 (Figure S1A), which results in a loop shift compared to PLD4 (Figure 4A). The extra clamp in PLD4 may confer stronger binding to the substrate and catalytic product, which is supported by the captured product only being observed in the PLD4 structure but not in PLD3. To assess if stronger substrate binding by the clamp may account for the lower activity of PLD4 on complex oligonucleotide substrates compared to PLD3, we mutated the clamp residue and found that PLD4 L183G had no enzymatic activity by our fluorophore-quencher assay, while PLD3 G170L exhibited even higher activity than WT. This surprising result suggests that other amino acids that differ in the G170/L183 loops of mPLD3 and hPLD4 (Figure S1A) may also contribute to the enzymatic activity. PLD3 cleaved both structurally diverse and homogeneous (oligo-dT) ssDNAs carrying a 5′-FAM group, albeit to a lesser extent, whereas PLD4 was not active (Figure 4C), suggesting that PLD4 could be more selective for substrates compared to PLD3. The extra clamp of PLD4 and attendant constriction of the substrate pocket may explain its selectivity (Figure 4A).

**Figure 4.**
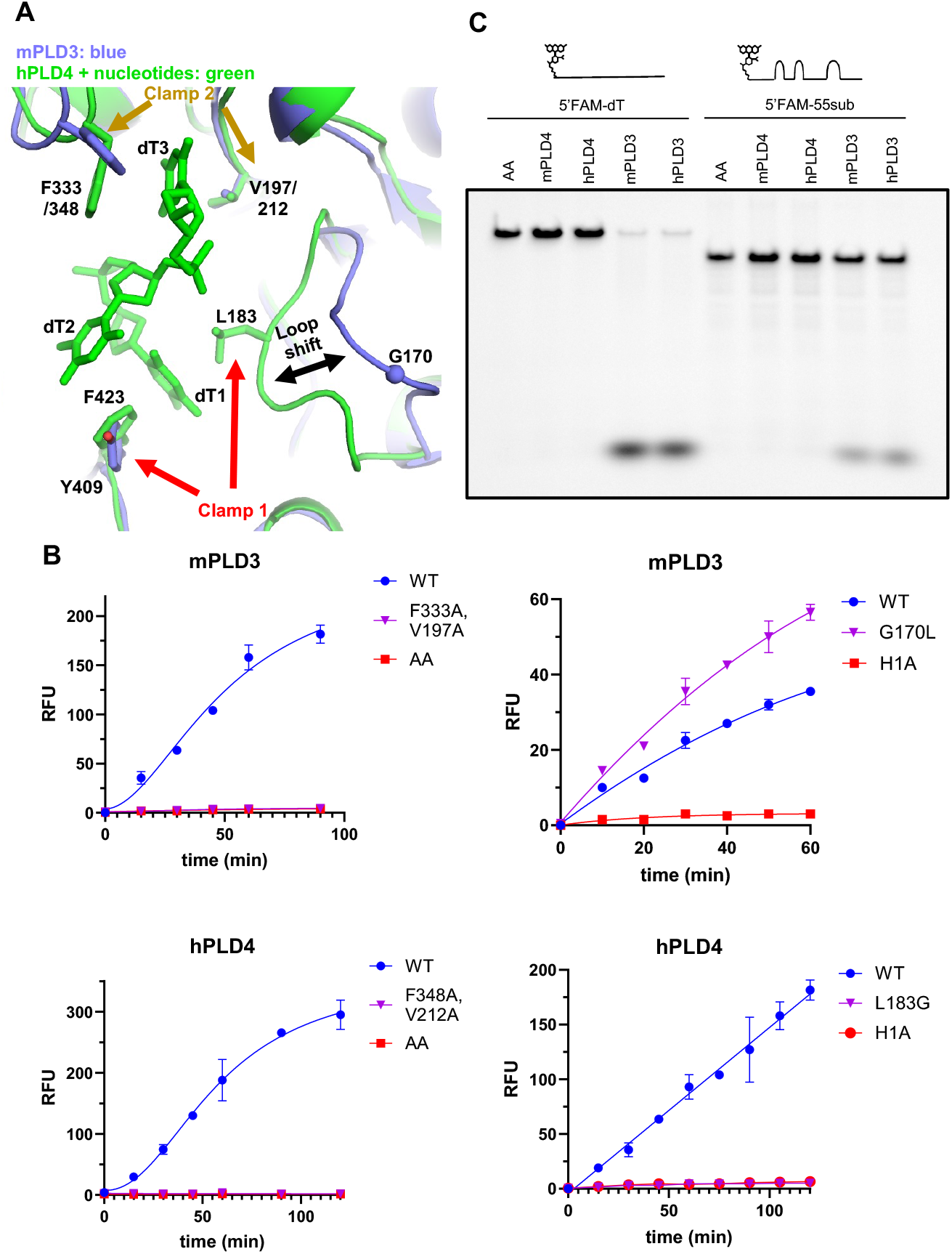
Structural comparison reveals additional substrate-binding hydrophobic clamp in PLD4. **(A)** Structural comparison between the active sites of mPLD3 (blue) and hPLD4 (green). Substrate-clamping amino acid residues are highlighted by arrows (see Figure S1A for details). **(B)** Mutational analysis of clamp 1 and 2 on activity of PLD3 and PLD4. Left two panels: mutation of key hydrophobic residues on clamp 2 abolishes enzyme activity of PLD3 and PLD4; Right two panels: Swapping the key residues of clamp 1 increases PLD3 activity (G170L) but decreases PLD4 activity (L183G). AA/H1A: both/first catalytic histidine(s) mutated to alanine. **(C)** Ability of PLD3 but not PLD4 to digest ssDNA substrates carrying a 5′-FAM motif, possibly because of a clash with the L183-containing loop in PLD4.

### Characterization of disease-associated PLD3/4 mutants

V232M is the first reported and most common (Figure S6) PLD3 variant genetically associated with late onset Alzheimer’s disease (AD) [41], although the association was questioned by other researchers [42–44]. Subsequently, I163M, R356H, and P410S were identified as risk variants for AD in a Chinese Han cohort[26,45,46]. L308P of hPLD3 has been genetically associated with SCA, and its 5¢-exonuclease activity was found to be impaired [47,48]. In a recent paper, it was suggested that PLD3 can degrade mitochondrial DNA to block cGAS/STING activation and thereby possibly prevent neurodegenerative diseases [49,50]. For PLD4, several disease-associated mutants have been reported in the Genome Aggregation Database (gnomAD) [51] (Figure S6). We expressed recombinant proteins of several of these variants carrying single point mutants in the luminal domain and characterized their enzyme activity and effect on TLR9 activation (Figure 5A-5B). For human PLD3, I163M partially lost exonuclease activity, and L308P almost completely lost enzymatic function. Their corresponding mutants in mouse PLD3, I163M, and L306P, had similarly reduced enzyme activities (Figure 5A-5B). Unexpectedly, hPLD3-V232M and the equivalent mutation mPLD3-V230M slightly increased enzyme activity. Size exclusion chromatography (SEC) of these mutants revealed that I163M, L308P of hPLD3 and R235Q, S283L of hPLD4 tended to form larger particles or aggregates, while V232M of hPLD3 only showed a slightly higher amount of larger particles (Figure 5C). We then measured thermostability of these mutants by differential scanning calorimetry (DSC). All PLD3/4 mutants had slightly reduced melting temperatures (Figure 5D). The SEC and thermostability results suggest that the disease-related mutations of PLD3 and PLD4 can destabilize the enzymes.

**Figure 5.**
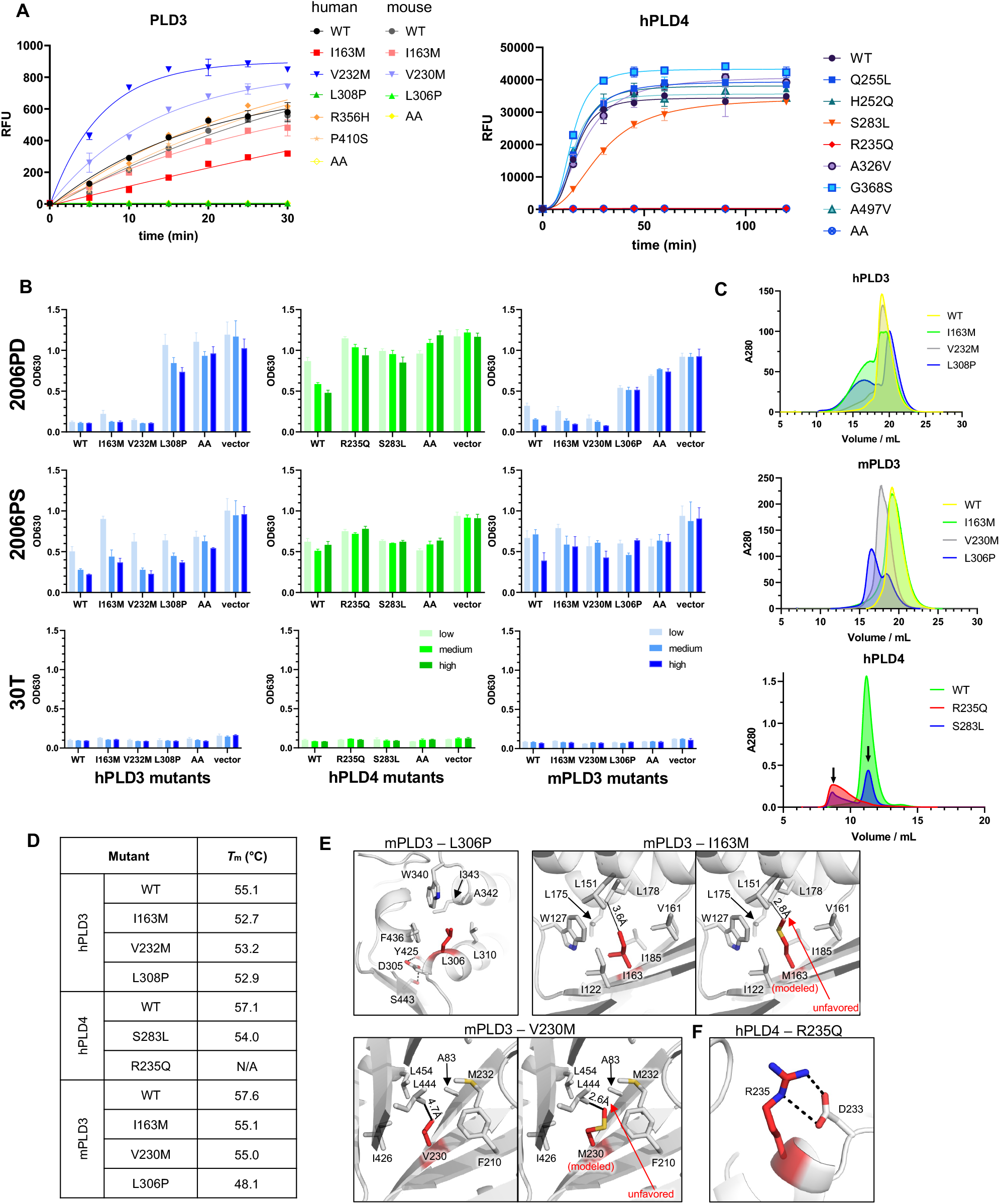
Functional characterization of variants of PLD3 and PLD4. **(A)** Analysis of activity of selected PLD3 and PLD4 missense mutant proteins by fluorophore-quencher assay. **(B)** Cell-based stimulation assay for selected PLD3 and PLD4 mutants. HEK293Blue^TM^ hTLR9 cells lacking PLD3 KO were reconstituted with different doses of the indicated PLD3 and PLD4 variants and stimulated with either 2006PD, 2006PS, or 30T (control). Readout reports NFκB activation. **(C-E)** Effect of the indicated point mutations of PLD3 and PLD4 on protein stability. **(C)** Size exclusion chromatography (SEC) of wile-type and mutated PLD3 and PLD4 with His-Myc tag. The aggregation peaks are represented by the early retention fractions. **(D)** Effects of selected PLD3 and PLD4 mutations on observed melting temperature. **(E)** Detailed structures of sites of relevant point mutations. mPLD3: L306, I163 and V230 are buried inside hydrophobic pockets of the protein; modeling of I163M and V230M mutants shows increased steric hindrance. **(F)** Location of PLD4 R235Q. Salt bridges are represented by black dashed lines. See also Figure S6 and Table S2.

We next modeled the mutations to our structures of PLD3 and PLD4 (Figure 5E-F). Mouse PLD3 L306 (corresponding to human L308) is located in the second position of an alpha helix consisting of eleven amino acids, where the side chain is stabilized by a hydrophobic core of PLD3. Mutating L306 to proline would disrupt the helix (Figure 5E). The side chain of mPLD3 I163 also falls into a hydrophobic core, where the closest residue L178 is 3.6-Å away from I163, which is well accommodated. However, the longer side chain of methionine is likely to be unfavored because of the shorter distance of only 2.8 Å to L178. Likewise, mutating V230 with a short side chain to methionine may cause unfavorable contact with the adjacent L444 (only 2.6 Å, Figure 5E). For human PLD4, R235 is stabilized by forming two salt bridges with D233 (Figure 5F), while the R235Q mutation would disrupt the salt bridges and potentially reduce the stability of hPLD4. We also calculated the free energy cost upon mutation using I-Mutant 3.0[52]. The results showed that the ΔΔG values are lower than −0.5 kcal/mol for all these three mutations, suggesting that the mutations would decrease the stability of PLD3 (Table S2). Intriguingly, our structural data showed that although almost all of the mutations are distant from the active sites (except for hPLD4-R235Q), they affect the enzyme activity and protein stability, which may explain their disease-association phenotypes.

## DISCUSSION

High-resolution structures of human PLD4 and mouse PLD3 in complex with substrates enabled us to deduce a possible catalytic mechanism of the two novel 5′-exonucleases and to explore structure-function relationships. Our data are consistent with a model in which PLD3 and PLD4 utilize a ping-pong mechanism characteristic of PLD family enzymes, where an active site histidine first attacks the phosphodiester bond, yielding covalent attachment to the phosphate, followed by release of the product facilitated through the second histidine [53]. PLD3 and PLD4 both cleave ssDNA and ssRNA substrates, but their distinct enzyme activities and substrate preferences may shape their biological functions. PLD3 and PLD4 are in part functionally redundant. While PLD3 is ubiquitously expressed in somatic tissues, PLD4 is limited to certain antigen presenting cells, especially dendritic cells. Here we show that the extra clamp PLD4 stabilizes the first base of the substrate as demonstrated in our crystal structures and may also affect product release. The structural and biochemical differences could help explain why PLD3 and PLD4 are both conserved for degradation of excess nucleic acids. We speculate that PLD3 is a general and more active ssDNA/RNA degrader, while PLD4 may be specialized in processing or protecting certain DNA/RNA sequences, thus regulating recognition by nucleic acid-sensing TLRs. PLD4’s lower enzyme activity and tunable expression may be important for dendritic cells to respond to pathogen stimulation in early viral infection. Since many chemical modifications have been identified on DNA and RNA molecules, it is also worth investigating the selectivity of PLD3/4 catalysis of nucleic acid substrates carrying methylation or other modifications, as well as the biological purpose of the selectivity. Alternatively, or in addition, as PLD4 has a relatively low pH optimum, its role may be to initially permit, then later to terminate signaling of endosomal TLRs by destroying their ligands as lysosomes acidify.

An unexpected finding of this study was that oligonucleotides carrying a 5′-phosphate were cleaved by PLD3 and PLD4, resulting in a phosphate covalently linked to position 3 (tau) of a histidine in the active site. These findings suggest that PLD3 and PLD4 can also act as 5′-polynucleotide phosphatases. Additionally, the enzyme activity of PLD3 and PLD4 is inhibited by nucleic acids carrying a 5′-phosphate. Here we illustrate the inhibitory mechanism of 5′-Pi nucleotides (Figure 2A). A previous study of the “spleen exonuclease” (in retrospect, almost certainly PLD3 [54]) found that an RNA substrate carrying a 5′-phosphate was cleaved by a two-step process involving a slow removal of elements at the 5′-end, leading to a 5′-OH form, followed by rapid exonuclease cleavage generating 3′-phosphates, though the authors proposed cleavage downstream of the second phosphate, not a dephosphorylation reaction, and the enzyme:substrate ratio was unknown [55]. Our data therefore indicate that both PLD3 and PLD4 harbor a 5′-phosphatase activity, albeit an inefficient one, as phosphate release appears to be slow. It is unclear if 5′-phosphorylated nucleic acid substrates are normally present in endolysosomes, as endonucleases in that compartment, such as DNase II and RNAse T2, cleave nucleic acids to yield fragments with 5′-OH groups. In addition, lysosomes carry two well-known phosphatases, acid phosphatase 2 (ACP2) and ACP5, with the ability to dephosphorylate mannose-6 phosphate modifications of proteins [56], and ACP2 has the ability to dephosphorylate AMP in acid conditions [57]. It will be interesting to investigate whether ACP2 and ACP5 also function as nucleotide phosphatases. It is perhaps significant that deficiency of these phosphatases is linked to inflammatory diseases [58,59], which we would predict arise from the resulting inhibition of PLD3 or PLD4 with attendant activation of nucleic acid sensors. In any case, some RNA virus genomes carry 5′-monophosphate [60], which may inhibit PLD3 and PLD4, possibly triggering downstream sensors for host innate immunity. Overall, PLD3 and PLD4 may be important drug targets. The structures of PLD3 and PLD4 along with their proposed mechanisms of catalysis should help aid efforts to design compound inhibitors [61].

### Limitations of the Study

Although we intended to generate structures of human PLD3 and PLD4, we were unable to obtain crystals of hPLD3. Fortunately, mPLD3 is 94% identical to hPLD3, so we believe the conclusions obtained are valid also for human. Human and mouse PLD3 appeared to have identical activities in all cases where they were compared, including both exonuclease and phosphatase assays.

Many PLD family members contain symmetric active sites where each domain consists of a histidine and a lysine, as well as an acidic residue that interacts with the histidine-N1 atom, e.g. *Streptomyces* PLD [53], Nuc [19], BfiI [20] (Figure S7). One of the acidic residues donates an electron to facilitate the nucleophilic attack of the nearby histidine, while the other acidic residue positions the vicinal histidine. In contrast, our structures showed that PLD3 and PLD4 are asymmetric in terms of the acidic residues—the first glutamic acid exhibits a nearly identical location as that of Nuc, but PLD3/4 lack the second acidic residue (Figure S7). Instead, a water molecule coordinates the side chain hydroxyls of T428, S429, as well as the backbone carbonyl oxygen of A440, and the H199-N1 atom to position the histidine in the active site. The different roles and functions of the symmetric and asymmetric or pseudosymmetric PLDs remain to be studied.

Similar to several previously reported nucleases [3], the exonuclease activity of PLD3 and PLD4 does not require divalent metal ions to facilitate the catalysis. However, we unexpectedly found that Fe^2+^, Cu^2+^, and Zn^2+^ inhibited the enzyme activity, as the divalent metals may be chelated by the two histidines at the active site [40] and compete with substrate binding. A similar phenomenon may happen in other enzymes with di-histidines at the active sites, e.g. human PLD1 [62], PLD2 [63], and Tdp1 [64]. In Alzheimer’s patients, it is reported that Fe^2+^ and Cu^2+^ levels are elevated [65,66]. Excess Fe^2+^ was also observed in the biological process of ferroptosis [67]. Elevated metal ion levels may then inhibit PLD3 activity and thereby boost inflammation, and reduce activity of other di-histidine-containing enzymes. The physiological relevance of the inhibitory effect of Fe^2+^, Cu^2+^, and Zn^2+^ is unclear.

## ACKNOWLEDGMENTS

We thank Robyn Stanfield, Xiaoping Dai, Yuanzi Hua, Henry Tien, Panpan Zhou and Wenli Yu for support and technical expertise. We thank Rajasree Kalagiri, Jia Ning, and Tony Hunter for the gift of anti-pHis antibodies and insightful discussions on phosphohistidines. We thank Jean-Laurent Casanova for genetic information and advice. This work was supported by R01AI142945 and RF1AG070775 to D. N. for studies on the biology of PLD4 and PLD3 and Skaggs Institute for Chemical Biology at Scripps Research (I.A.W.). X-ray data sets were collected at the GM/CA@APS-23ID-B and 23ID-D beamlines, which have been funded in whole or in part with Federal funds from the National Cancer Institute (ACB-12002) and the National Institute of General Medical Sciences (AGM-12006). We thank the Burton lab for assistance with protein expression and purification. This research used resources of the Advanced Photon Source (APS), a U.S. Department of Energy (DOE) Office of Science User Facility operated for the DOE Office of Science by Argonne National Laboratory under Contract No. DE-AC02-06CH11357. The contents of this publication are solely the responsibility of the authors and do not necessarily represent the official views of NIH or the US Government. The funders had no role in study design, data collection and analysis, decision to publish, or preparation of the manuscript.

## AUTHOR CONTRIBUTIONS

M.Y., L.P., D.H., D.N. and I.A.W. conceived the study. M.Y. determined the crystal structures. M.Y., Z.F. and X.Z. conducted structural analysis. L.P., D.H., F.L., A.G. J.T., J.M. performed subsequent biochemical experiments. L.P., M.Y., I.A.W., and D.N. wrote the manuscript; all authors reviewed and revised the paper. The funding was secured by D.N. and I.A.W.

## METHODS

### Establishment of constructs of PLD3 and PLD4

PLD3 and PLD4 recombinant protein constructs were cloned into a SuExp vector and their sequences can be found on Addgene (#173851-173853, His-Myc tagged; #201243-201245, full length PLD). Single point mutant plasmids were constructed using Q5^®^ site-directed mutagenesis kit (NEB, E0554S) according to manufacturer’s instructions. Lentiviral related PLD3/4 plasmids were cloned into pBOBI vector that has a C-terminal FLAG tag after digestion by restriction enzymes BamHI and XhoI (NEB R3136S, R0146S). All these plasmids were prepared with endotoxin-free kit (Takara 740426) and sterile filtered before transfection.

### Lentivirus production

293T cells were seeded into 6 well plates a day prior to transfection. 2 μg pBOBI-PLD3/4, 1.5 μg dR8.2 (Addgene #8455) and 0.5 μg pMD2.G (Addgene #12259) were transfected with Lipofectamine 2000 (11668027, ThermoFisher) according to manufacturer’s instructions. 12-16 h after transfection, 5 mL fresh DMEM medium was added, and the supernatants were collected and filtered.

### Protein expression

Recombinant PLD3 and PLD4 proteins were expressed in Expi293F cell line according to the manufacturer’s instructions (ThermoFisher). In brief, 24 μg plasmids and 24 μL FectoPro transfection reagent were mixed in 3 mL Opti-MEM medium (ThermoFisher), then incubated for 20 min at RT, and added to 30 mL 3×10^6^/mL Expi293 cells in 293Expi medium. 0.27 mL 45% D-α-glucose and 0.3 mL 300 mM valproic acid (VPA) were added to cell culture 24 h after transfection. The reactions were scaled up proportionally. For crystallization, HEK 293S cells were used to express N-terminally His_6_-tagged luminal domains of mPLD3 (Uniprot ID O35405, residues 63-488) and hPLD4 (Uniprot ID Q96BZ4, residues 60-506). In brief, 400 μg of each plasmid was mixed with 1 mL 10 g/L polyethylenimine (PEI) in 50 mL Opti-MEM, incubated 20 min at RT and added to 500 mL 3×10^6^/mL HEK 293S cells culture in Freestyle 293 expression medium (ThermoFisher). The cells were cultured for 5 days, centrifuged and supernatants were filtered through 0.22 μM filter. One milliliter of Ni-NTA agarose beads were added to the supernatants and incubated with gentle shaking at 4°C overnight. The beads were filtered out, washed with 20 mL of 20 mM imidazole in PBS, and eluted with 5 mL of 500 mM imidazole in PBS, followed by size exclusion chromatography, and buffer exchange into acetate buffer (20 mM sodium acetate, 125 mM NaCl, pH 6.0). The proteins were then ultrafiltered with 30 kDa centrifugal filter units (MilliporeSigma). Protein concentrations were assessed by absorbance at A280 using a nanodrop device with the extinction coefficient calculated by Expasy (https://web.expasy.org/protparam/).

### Reporter cell lines

All 293 reporter cell lines in this paper were cultured in DMEM supplied with 10% FBS, 1% penicillin/streptomycin and 2 mM glutaMAX (Thermo Fisher), unless specified otherwise. HEK293Blue^TM^ hTLR9 cell line was purchased from InvivoGen and cultured according to manufacturer’s instructions. PLD3 KO cell line was generated from HEK293Blue^TM^ hTLR9 by transfecting 2 μg hSpCas9-sgRNA expressing plasmid (Addgene #99154) cloned with gRNA sequence 5′-guccucauucuggcgguugu-3′. After 16 h, mCherry^+^ single cells were sorted into 96-well plates and cultured for 4 weeks. Single clone cells were harvested, genotyped by PCR and Sanger sequencing and one clone with both PLD3 alleles frameshifted was obtained. PLD3 KO-HEK293Blue^TM^ hTLR9 cells were infected with lentivirus to generate cell lines that stably express full length PLD3, PLD4 or the corresponding mutants. Different amounts of lentivirus were added to adjust the expression dose of PLD3 or PLD4. RT-PCR and western-blot were performed to determine the relative expression level of each cell line.

### PLD3 and PLD4 enzyme assays

For cell-based enzymatic activity assay, PLD3^-/-^ HEK293Blue^TM^ hTLR9 reporter cell line was reconstituted with wild-type or variant alleles of PLD3 or PLD4. Cells were seeded into 96-well plates (80,000 per well) and 1 μM CpG-containing oligodeoxynucleotide (ODN) TLR9 agonists 2006-PD and 2006-PS (5’-tcgtcgttttgtcgttttgtcgtt-3′) carrying phosphodiester (PD) or phosphorothioate linkages (PS) or control ODN containing 30 thymidines (30dT) was added to the medium [68]. After culturing for 20 h, 10 μL supernatants were collected from each well and added to 90 μL QUANTI-Blue substrate solution (InvivoGen) to detect expression of secreted embryonic alkaline phosphatase (SEAP) under NF-κB regulatory elements. The reactions were incubated at 37°C for 30 min, and optical density at 630 nm (OD_630_) was measured with a plate reader. PLD3 and PLD4 efficiently digest 2006-PD but not 2006-PS in vivo to suppress TLR9-driven NF-κB signaling.

Both gel-based and fluorophore-quencher assays were used to measure the activity of PLD3 and PLD4. For the gel-based assay, the 5′-FAM dT contains a 50 T oligonucleotide, and the sequence of the 5′-FAM 55SUB is 5′-TCCATGACGTTCCTGATGCTAAGTATGCACTTCATCGTCAAGCAATGCTATGCA. 20 nM PLD3 and PLD4 were incubated with substrates at 2 µM final in acetate buffer (50 mM acetate and 20 mM NaCl, pH 5.6 and 4.4, respectively) for 2 hours at 37°C. 20% TBE-PAGE gel was used to separate the ssDNA. For the fluorophore-quencher assay, the reaction condition for PLD3 was 2 nM enzyme in pH 5.5 MES buffer (50 mM MES-HCl, 100 mM NaCl, 10 μg/mL OVA), room temperature; for PLD4 100 nM enzyme in pH 4.7 acetate buffer (50 mM NaAc-HAc, 100 mM NaCl, 10 μg/mL OVA), 37°C. The final concentration of the fluorophore-linked substrate dT_5_ or 55nt oligodeoxynucleotide (ODN) was 2 μM. The fluorophore-quencher assay was carried out using black 384-well plates (Greiner, REF 788076) with a total volume of 10 μL. For normal assay endpoint, incubation time of PLD3 was 45 min; PLD4 2 h. 5 μL 1 M Tris-HCl buffer, pH 8.8 was added to quench the reaction, and fluorescence signal was quantified on a plate reader.

### Dinucleotide substrate assay

Adenosine deaminase (ADA) was purchased from Worthington Biochemical Corp and was resuspended in 1 mL water, then was diluted to 20 μg/mL in 0.5x PBS. The dinucleotide substrate UpA was purchased from TriLink Biotechnologies. The final reaction buffer was 50 mM phosphate and 20 mM NaCl, pH 7.5. Each reaction consisted of a total volume of 250 μl with dinucleotide substrate (40 μM), ADA enzyme (2 μg/mL) and one of the following: PLD3 (25 nM) or PLD4 (25 nM) or His to Ala PLD mutants (25 nM). The ADA only control had PBS added instead of PLD3/4 enzyme. The reaction was performed with the Nanodrop 3000 C in a heated quartz cuvette (37°C), and absorbance at 265 nm was measured every 10 s. The phospholipase to be tested was added after four initial absorbance readings (45 sec mark), and absorbance measured for another 4 min.

### Crystallization and structural determination

mPLD3 and hPLD4 were screened for crystallization using the 384 conditions of the JCSG Core Suite (Qiagen) on our robotic CrystalMation system (Rigaku) at Scripps Research. Crystallization trials were set up by the vapor diffusion method in sitting drops containing 0.1 μl of protein and 0.1 μl of reservoir solution. For the mPLD3 apo protein (13 mg/ml), crystals were grown in drops containing 12% PEG 3350, 0.5 mM MgCl_2_, 0.133 M di-ammonium tartrate, pH 6.6 and 15% (w/v) ethylene glycol at 4°C. Crystals appeared at day 7 and were allowed to grow for 30 days before mounting. To investigate the structural basis of mPLD3 catalysis, 13 mg/ml mPLD3 was crystallized in the presence of 5′Pi-(dT)_5_ (2-fold, molar ratio). Crystals were grown in drops containing 1.6 M (NH_4_)_2_SO_4_, 0.5 mM MgCl_2_, 0.1 M citric acid, pH 4.0 at 4°C. Crystals appeared at day 3 and were allowed to grow for 9 days and 30 days before mounting. We also co-crystallized 10 mg/ml hPLD4 with 5′Pi-(dT)_5_ (2-fold, molar ratio) in drops containing 0.2 M NaCl, 0.5 mM MgCl_2_, 0.1 M phosphate-citrate pH 4.2, and 10% (w/v) PEG3000 at 4°C. Crystals appeared at day 7, and were allowed to grow for 30 days. Crystals were harvested by soaking in reservoir solution supplemented with 15% ethylene glycol (w/v) as cryoprotectant. Diffraction data were collected at cryogenic temperature (100 K) at beamline 23-ID-B or 23-ID-D of the Advanced Photon Source (APS) at Argonne National Labs. Diffraction data were processed with HKL2000 [52]. Structures were solved by molecular replacement using PHASER [69]. Iterative model building and refinement were carried out in COOT [70] and PHENIX [71], respectively.

### Analysis of phosphate transfer from 5′-phosphorylated DNA to PLD3 and PLD4

Seventy picomoles of oligonucleotide (40T) was labelled with polynucleotide kinase (PNK) in PNK buffer (NEB) in the presence of *γ*^32^P-ATP (Perkin-Elmer) at 37°C for 30 minutes. The reaction was stopped by heat inactivation at 75°C for 10 minutes. Free *γ*^32^P-ATP was removed by size exclusion chromatography over G-25 microspin column (GE illustra). Purified recombinant PLD proteins (42 pmol) were incubated with 7 pmol of 5′-labelled oligonucleotide for 1 hour in 20 mM NaCl 50 mM acetate buffer (pH 5.2) before SDS sample buffer was added and the proteins heated to 75°C for 10 minutes. Proteins were electrophoresed on 4-12% SDS NuPAGE gels (Thermo Fisher Scientific) prior to being transferred to PVDF membranes. The presence of radio-labelled proteins on the membrane were revealed by autoradiography. The equivalent amount of protein from each reaction was also electrophoresed on 4-12% SDS NuPage gels and detected by Simply Blue Safe stain (Invitrogen).

### Phospho-histidine western blot assay

Recombinant PLD3 and its variants (1 μM) were incubated for 2 h with oligos carrying 5′-phosphate or 5′-hydroxyl group in 50 mM 2-(N-morpholino)ethanesulfonic acid (MES) pH 6.5, 125 mM NaCl. The reaction was quenched with 1 M of Tris-HCl (pH 9.0) and subjected to western blotting analysis. The proteins were isolated with pH 8.8 Tris-PAGE gel at 4°C, transferred to PVDF blot and developed with anti-1-pHis (sc-1-1) or 3-pHis (sc-39-4) antibody [34].

### Size exclusion chromatography (SEC) and differential scanning calorimetry (DSC)

Purified His-Myc recombinant PLD proteins were loaded to SEC (AKTA) for separation. The column used was Superdex 200 Increase 10/300GL, run at 0.75 mL/min with PBS buffer.

Fractions corresponding to WT PLD3 or PLD4 proteins were collected, concentrated with centrifugal filters and measured with DSC.

