## Supplemental Materials for "Structural and mechanistic insights into disease-associated endolysosomal exonucleases PLD3 and PLD4"

### SUPPLEMENTAL INFORMATION

A

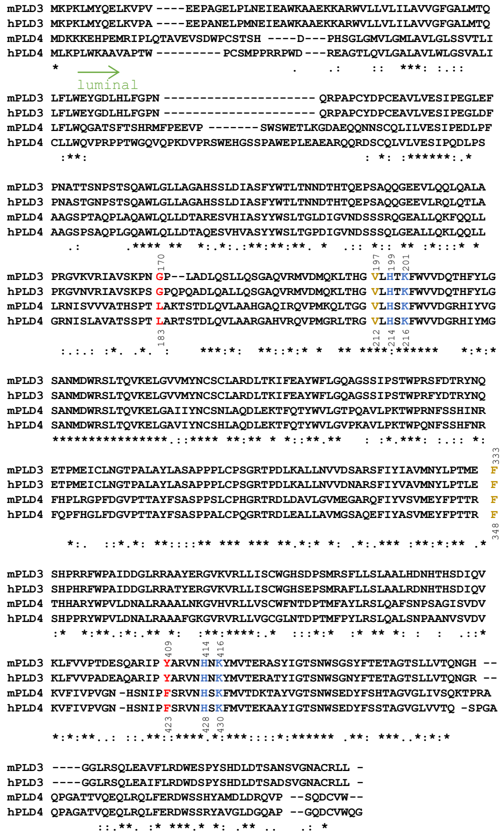

B

| | mPLD3 | hPLD3 | mPLD4 | hPLD4 |
|---|---|---|---|---|
| mPLD3 | 100 | 93.7 | 43.8 | 45.6 |
| hPLD3 | 93.7 | 100 | 43.2 | 44.3 |
| mPLD4 | 43.8 | 43.2 | 100 | 75.9 |
| hPLD4 | 45.6 | 44.3 | 75.9 | 100 |

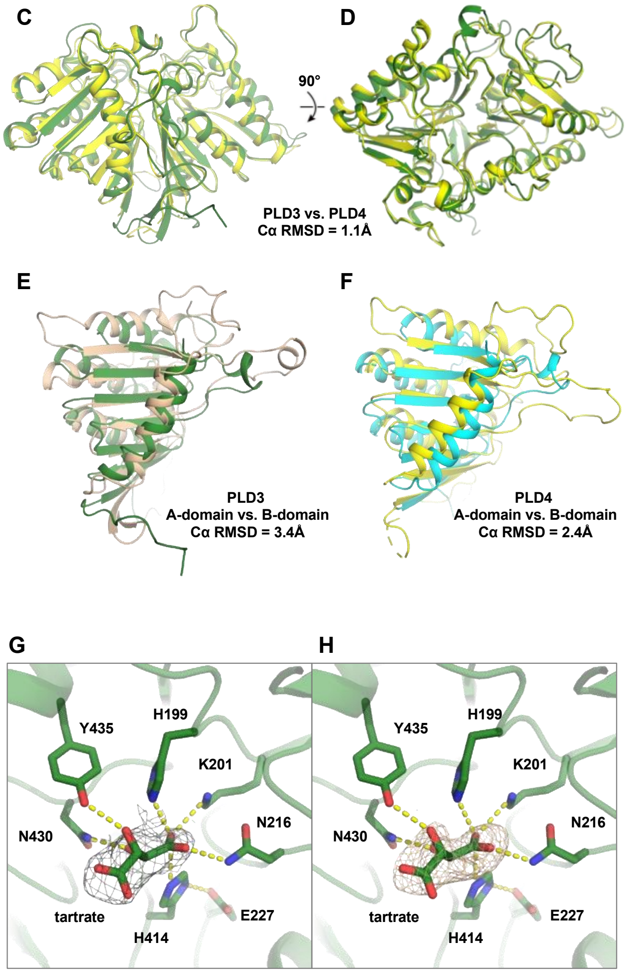


**Figure S1. Sequence and structure comparison of PLD3 and PLD4. (A)** Sequence alignment of PLD3 and PLD4. Mouse PLD3 (Uniprot No.: O35405), human PLD3 (Uniprot No.: Q8IV08), mouse PLD4 (Uniprot No.: Q8BG07), and human PLD4 (Uniprot No.: Q96BZ4). Residues conserved in all aligned PLDs are labeled by an asterisk (*), while a colon (:) and a period (.) indicate strongly similar and weakly similar sequences, respectively. Residues involved in phosphodiester-bond cleavage are highlighted in blue. Letters shown in red and brown represent residues that interact with the substrate DNA's first nucleotide, and its third nucleotide, respectively. Residue numbers that correspond to mPLD3 and hPLD4 are shown above and below the sequence alignment, respectively. The sequence alignment was performed with Clustal

Omega [1]. The luminal region of the PLDs is labeled with a green arrow. **(B)** Percent identity matrix for the sequence alignment. **(C-D)** Structural alignment between the crystal structures of **(C)** mPLD3 and **(D)** hPLD4. **(E)** Structural alignment between A-domain and B-domain of mPLD3. **(F)** Structural alignment between A-domain and B-domain of hPLD4. The structural alignments were carried out with PyMOL without further refinement cycles. **(G-H)** In the active site of mPLD3, a tartrate molecule from the crystallization solution was observed. Hydrogen bonds and salt bridges are represented by yellow dashed lines. **(G)** A $2F_o$-$F_c$ electron density map of the tartrate molecule is represented in a gray mesh contoured at 0.8σ. **(H)** An $F_o$-$F_c$ unbiased omit electron density map of the tartrate molecule is represented in a gray mesh contoured at 2.3σ.

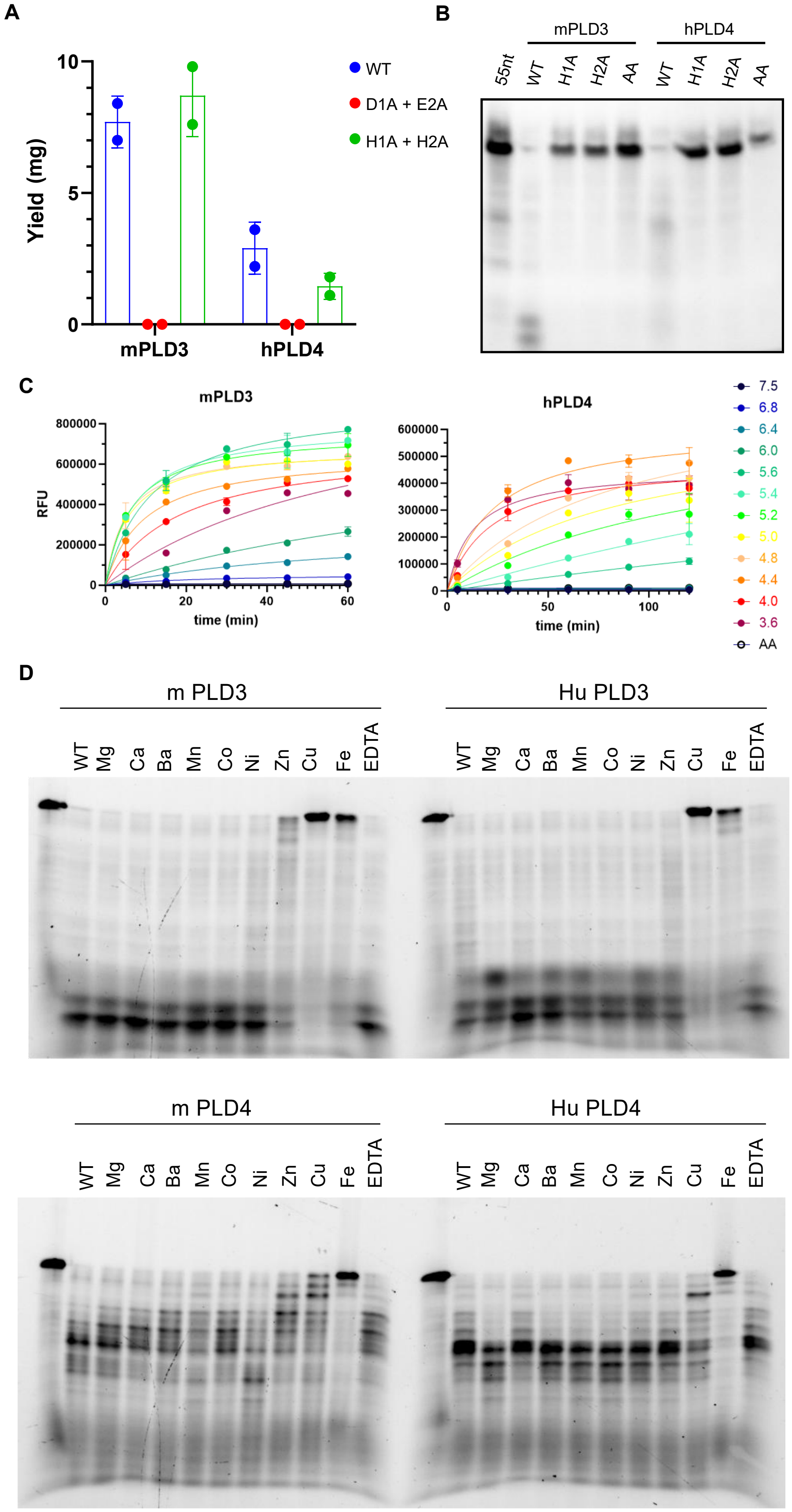

A
Yield (mg)
10
5
0
mPLD3
hPLD4
WT
D1A + E2A
H1A + H2A
B
mPLD3
hPLD4
55nt
WT
H1A
H2A
AA
WT
H1A
H2A
AA
C
mPLD3
RFU
time (min)
hPLD4
time (min)
7.5
6.8
6.4
6.0
5.6
5.4
5.2
5.0
4.8
4.4
4.0
3.6
AA
D
m PLD3
Hu PLD3
WT
Mg
Ca
Ba
Mn
Co
Ni
Zn
Cu
Fe
EDTA
m PLD4
Hu PLD4

**Figure S2**. **Effects of mutations and cation metals on PLD3 and PLD4. (A)** Expression yield for recombinant mPLD3 and hPLD4 by Expi293 cells per 30 mL medium. D1A + E2A: D206A + E421A (mPLD3) or D221A + E435A (hPLD4); H1A + H2A: both catalytic His mutated to Ala. Each red dot represents a data point. **(B)** Requirement of His residues in HKD motif for the PLD catalytic activity. H1A: first His was mutated to Ala; H2A: second His to Ala; AA: and both His to Ala. Reaction condition: 10 nM PLD3 or 100 nM PLD4, 2 μM 55nt substrate (55SUB 5′-TCCATGACGTTCCTGATGCTAAGTATGCACTTCATCGTCAAGCAATGCTATGCA-3′), reaction at 37°C for 2 h. **(C)** Exonuclease activity of human PLD4 and PLD3 at different pH values, as measured by a fluorophore-quencher assay. AA was used as a non-catalytic control. **(D)** Gel assay showing the inhibition of PLD3 and PLD4 by selected divalent cations. PLD proteins were dialyzed with 10 mM EDTA overnight, then against 1000-fold volume of PBS to remove any cations acquired from purification/elution. Reaction conditions were as follows. PLD4: 50 mM in NaAc buffer, pH 4.4, enzyme:substrate 1:100 (molar), reaction at 37°C for 2 h. PLD3: 50 mM in MES buffer, pH 5.6, enzyme:substrate 1:40, reaction at 37℃ for 1.5 h. In all reactions, 2 μM of 55Sub-FAM, and 2 mM of cations or EDTA was used.

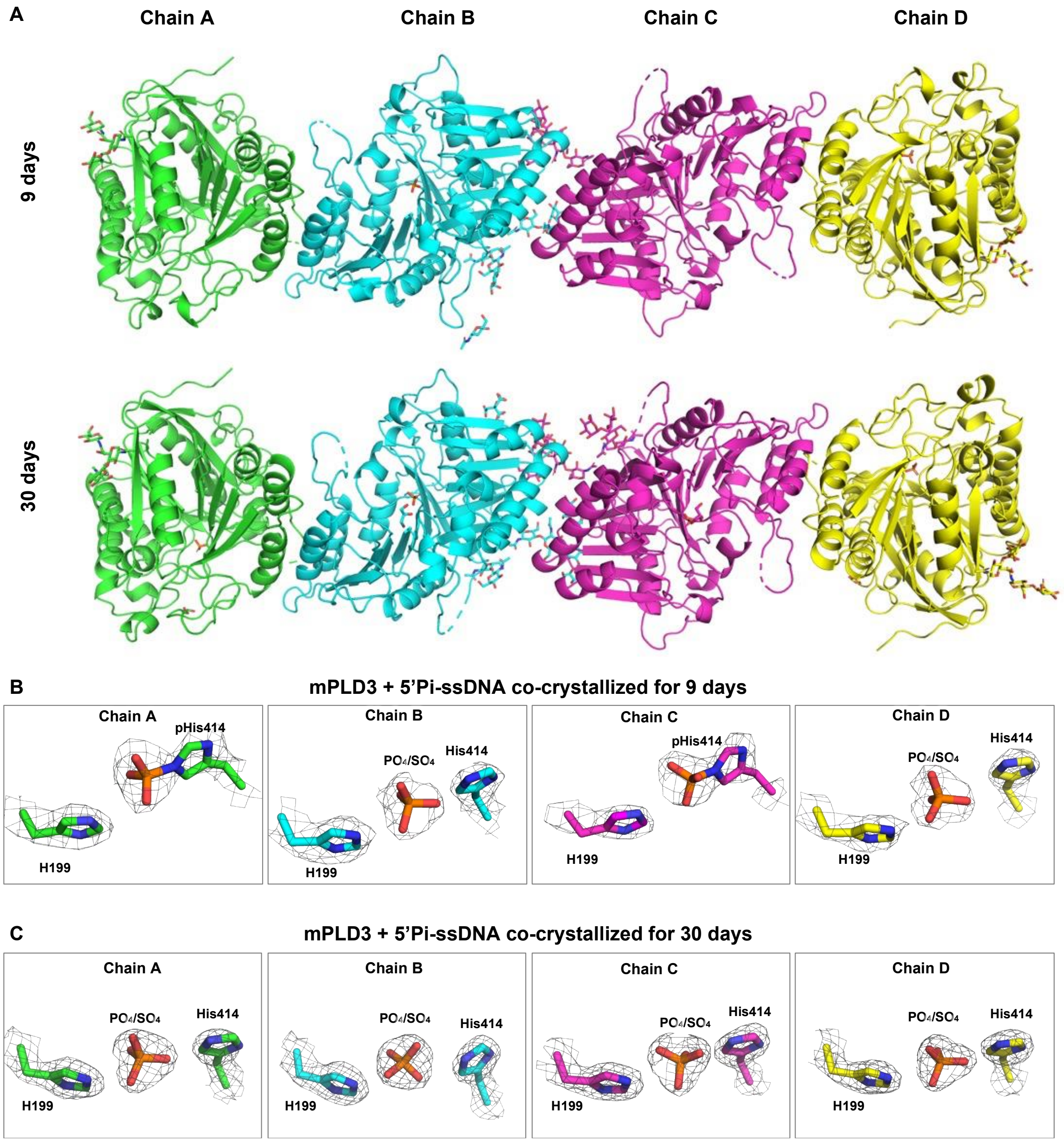


**Figure S3. Overall and active site structures of mPLD3 co-crystallized with 5′-Pi-ssDNA. (A)** Each asymmetric unit in the structures of mPLD3 co-crystallized with 5′-Pi-ssDNA for 9 and 30 days both contain four PLD3 molecules. **(B-C)** Active sites of each chain in the structures of mPLD3 co-crystallized with 5′-Pi-ssDNA for **(B)** 9 days and **(C)** 30 days. The $2F_o$-$F_c$ electron density maps are represented in a gray mesh contoured at 1.5σ.

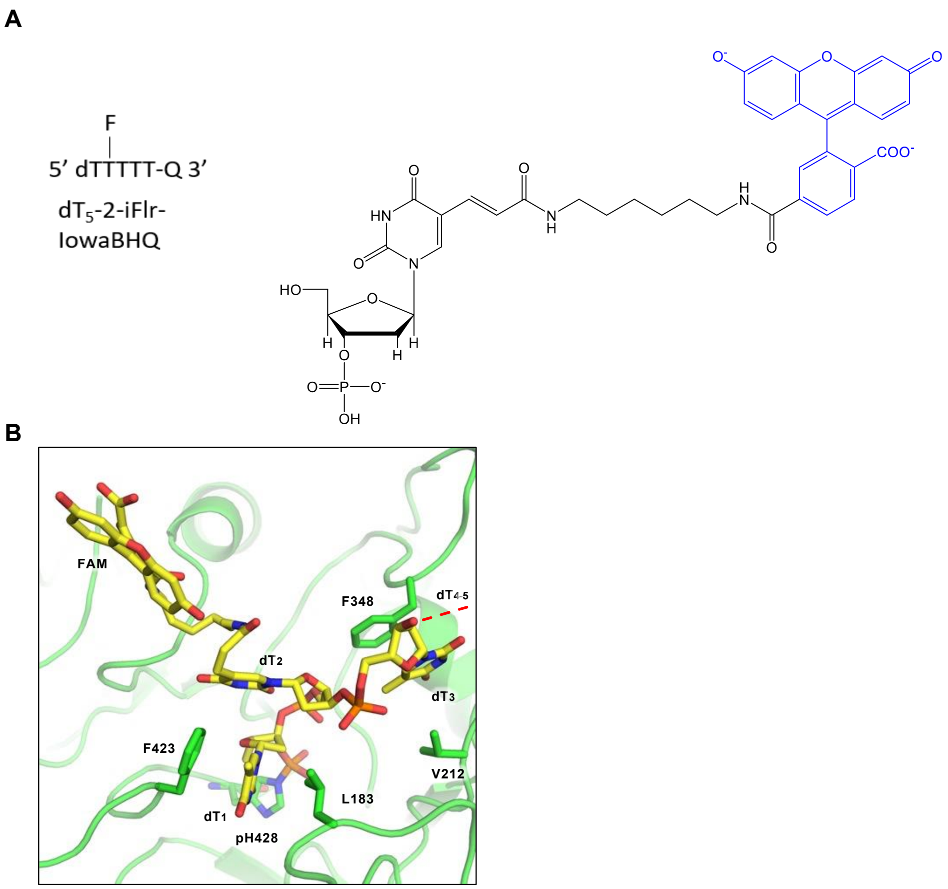


**Figure S4. Modeled structure of nucleotide-linked fluorescein of the fluorogenic substrate in the PLD4 active site. (A)** Chemical structure of the fluorophore-quencher substrate (blue) and details of the thymidine-linked FAM (abbreviated as 'F'). **(B)** A modeled structure of hPLD4 bound to substrate with thymidine-linked FAM. Red dashed lines represents the unmodeled $dT_4$ and $dT_5$.

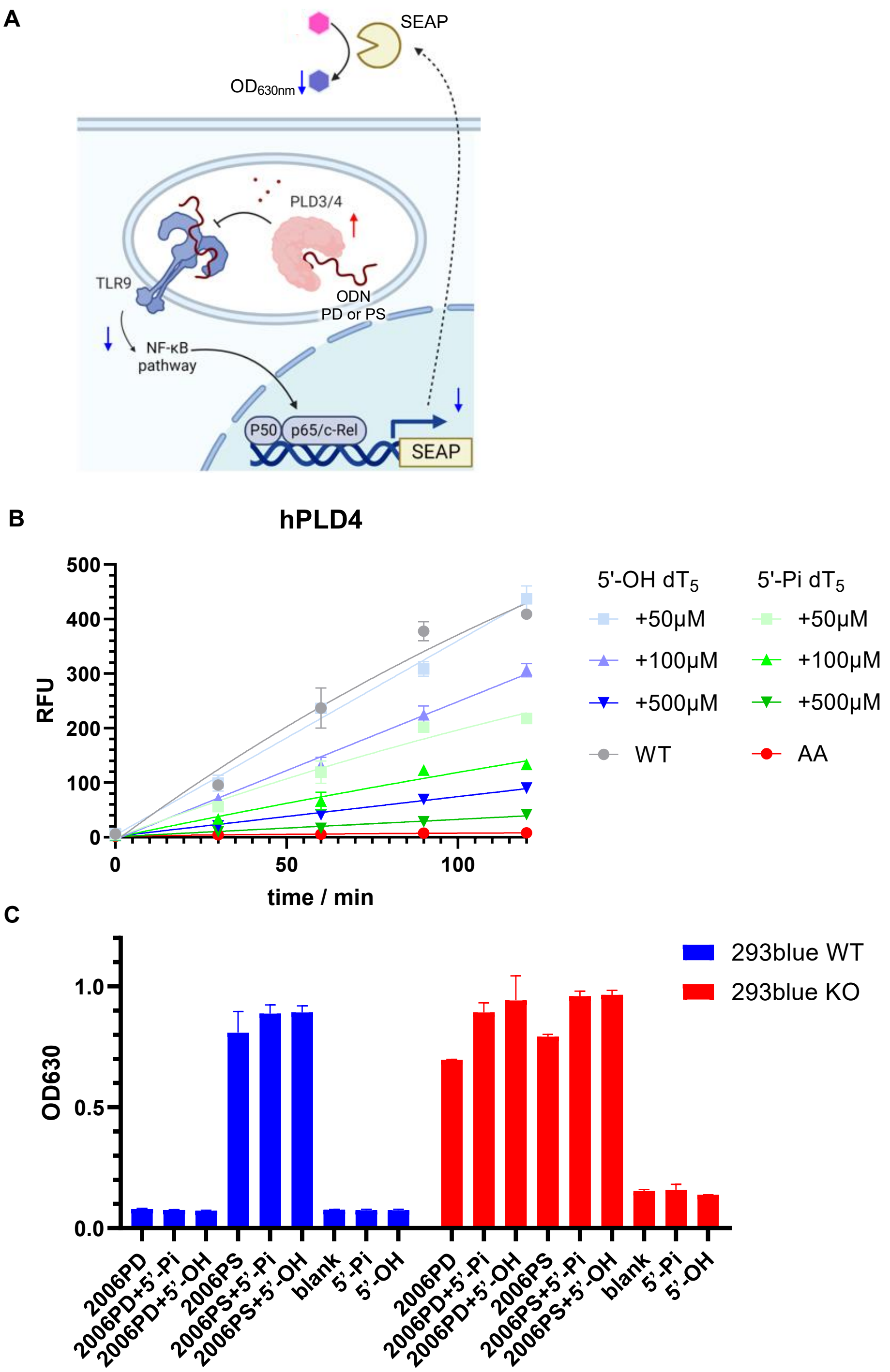


**Figure S5. Analysis of inhibitory effect of 5′-phosphorylated oligonucleotides in an in vitro enzyme assay or bioassay. (A)** Scheme of the cell-based assay for PLD3/4 enzyme activity. HEK293Blue$^{TM}$ hTLR9 reporter cells can be stimulated by TLR9 agonists (ODNs including

2006PD and 2006PS, represented by brown lines), thereby activating downstream NF-κB pathway signaling. Secreted embryonic alkaline phosphatase (SEAP) is produced under the control of NF-κB promoter. SEAP is secreted to supernatant and catalyzes the conversion of a colorimetric substrate from pink to blue, which can be quantified by absorbance at the wavelength of 630 nm ($OD_{630}$). The reporter cells with PLD3 knockout (KO) were transfected with PLD3/4 to digest ssDNA (here 2006PD), thus reducing TLR9-driven NF-κB reporter signaling. The higher the exonuclease activity (represented by a red up arrow), the less the $OD_{630}$ will be measured (blue down arrows). The figure was created by the BioRender software. Details of this assay were reported in our previous study [2]. **(B)** Analysis of dose-dependent inhibition of PLD4 by either 5′-Pi-$dT_5$ or 5′-OH-$dT_5$. In brief, 100 nM PLD4 and 2 μM iFr-5-dT were mixed in NaAc reaction buffer in the presence of escalating doses of 5′-Pi-$dT_5$ or 5′-OH-$dT_5$. The reactions were quenched at different time points with 1 M Tris, and the fluorescent signal was measured. **(C)** Cell experiment showing excess 5′-Pi-$dT_5$ was unable to inhibit PLD3 activity as measured by HEK293Blue$^{TM}$ hTLR9 reporter cell line. The cells (WT or PLD3 KO) were stimulated with 1 μM 2006PD in the presence of 1 mM 5′-OH-$dT_5$ or 5′-Pi-$dT_5$. No significant inhibition of WT PLD3 activity was observed. PLD4 efficiently digested oligonucleotide substrates that contains phosphodiester (PD) linkages but not phosphorothioate linkages (PS). 5′-OH = 5′-OH-$dT_5$; 5′-Pi = 5′-Pi-$dT_5$.

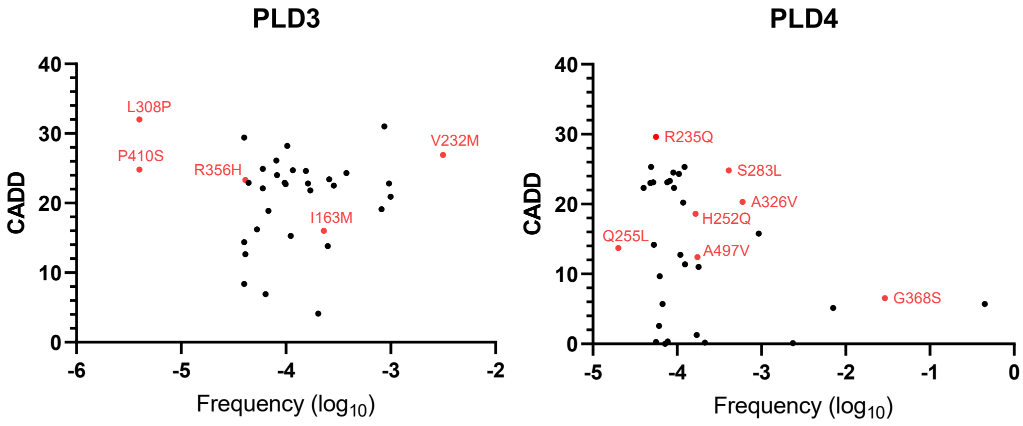


**Figure S6. Scatter plots of PLD3 and PLD4 disease-associated mutants.** Variant data were obtained from https://hgidsoft.rockefeller.edu/PopViz/. The plot indicates the selected missense mutations analyzed in this study (red). Y axis: CADD, Combined Annotation-Dependent Depletion. X axis: allele frequency.

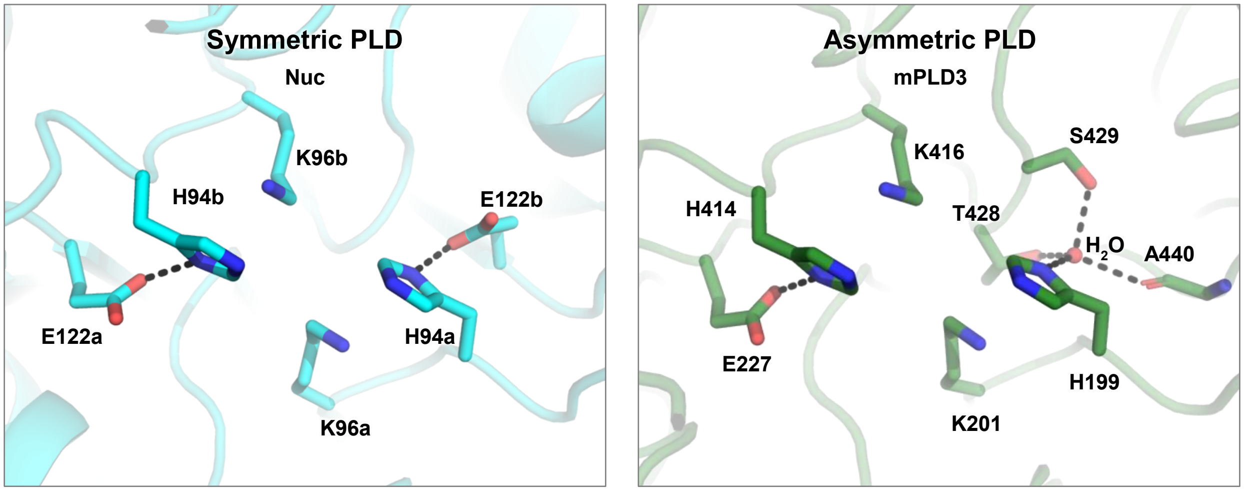


**Figure S7. Structural comparison of the active sites of symmetrical and asymmetrical PLDs.** Crystal structure of a phospholipase D family member, Nuc from *Salmonella typhimurium* (PDB 1BYS), is used to represent symmetric PLDs. Crystal structure of mPLD3 without nucleic acid substrate from this study is used to represent asymmetric (or pseudosymmetric) PLDs. Hydrogen bonds and salt bridges are represented by black dashed lines. Nuc is an interchain dimer where the residues in the first chain are labeled as 'a' and those in the second chain as 'b'. mPLD3 is an intrachain dimer, where all residues are labeled sequentially.

### Table S1. X-ray data collection and refinement statistics

| **Data collection** | | | | |
|---|---|---|---|---|
| | mPLD3 apo | mPLD3+5'Pi-$(dT)_5$ (9 days) [a] | mPLD3+5'Pi-$(dT)_5$ (30 days) [a] | hPLD4+5'Pi-$(dT)_5$ (14 days) [a] |
| Beamline | APS23ID-D | APS23ID-D | APS23ID-B | APS23ID-D |
| Wavelength (Å) | 1.0332 | 1.0332 | 1.0332 | 1.0332 |
| Space group | P $3_2$ 2 1 | P 1 | P 1 | P $3_1$ 2 1 |
| Unit cell parameters | | | | |
| a, b, c (Å) | 94.2, 94.2, 109.4 | 54.2, 54.2, 202.8 | 54.5, 54.8, 203.6 | 89.0, 89.0, 274.0 |
| α, β, γ (°) | 90, 90, 120 | 96.6, 89.2, 90.1 | 83.7, 89.1, 89.9 | 90, 90, 120 |
| Resolution (Å) [b] | 50.0-2.08 (2.12-2.08) | 50.0-2.75 (2.80-2.75) | 50.0-2.00 (2.03-2.00) | 50.0-3.00 (3.05-3.00) |
| Unique reflections [b] | 34,128 (3,344) | 57,208 (2,215) | 131,976 (10,415) | 26,067 (2,549) |
| Redundancy [b] | 18.7 (13.0) | 3.0 (1.9) | 2.3 (1.9) | 18.3 (13.7) |
| Completeness (%) [b] | 100 (99.8) | 95.5 (75.9) | 83.1 (69.3) | 99.8 (99.5) |
| $\langle I/\sigma_I \rangle$ [b] | 36.9 (2.1) | 11.9 (2.1) | 11.3 (1.0) | 27.9 (1.0) |
| $R_{sym}$[c] (%) [b] | 6.9 (64.4) | 12.1 (42.4) | 9.1 (70.1) | 9.6 (41.8) |
| $R_{pim}$ (%) [b] | 1.6 (17.6) | 8.0 (34.0) | 6.9 (58.7) | 2.3 (45.9) |
| $CC_{1/2}$[d] (%) [b] | 99.2 (94.0) | 96.7 (59.2) | 95.6 (52.9) | 99.6 (63.9) |
| **Refinement statistics** | | | | |
| Resolution (Å) | 45.4-2.08 | 48.0-2.75 | 37.1-2.00 | 44.7-3.00 |
| Reflections (work) | 34,105 | 57,195 | 131,924 | 26,067 |
| Reflections (test) | 1,679 | 4,069 | 6,468 | 2,549 |
| $R_{cryst}$[e] / $R_{free}$[f] (%) | 20.6/24.2 | 25.4/29.3 | 22.4/26.4 | 26.3/31.4 |
| No. non-H atoms | 3,747 | 13,353 | 13,744 | 6,286 |
| Macromolecules [g] | 3,515 | 13,158 | 13,125 | 6,172 |
| Ligands [h] | 26 | 10 | 72 | 114 |
| Solvent | 199 | 185 | 553 | 0 |
| Average *B*-values ($Å^2$) | 54 | 44 | 40 | 122 |
| Macromolecules [g] | 55 | 44 | 40 | 121 |
| Ligands [h] | 63 | 40 | 48 | 178 |
| Solvent | 49 | 34 | 39 | N/A |
| Wilson *B*-value ($Å^2$) | 39 | 37 | 33 | 110 |
| **RMSD from ideal geometry** | | | | |
| Bond length (Å) | 0.003 | 0.003 | 0.017 | 0.005 |
| Bond angle (°) | 0.65 | 0.65 | 1.8 | 1.0 |
| **Ramachandran statistics (%)** | | | | |
| Favored | 96.2 | 96.7 | 96.9 | 92.6 |
| Outliers | 0.00 | 0.00 | 0.25 | 0.26 |
| **PDB code** | 8V05 | 8V06 | 8V07 | 8V08 |

[a] The reaction in the crystal was allowed to proceed for different times before mounting and cryoprotection.
[b] Numbers in parentheses refer to the highest resolution shell.
[c] $R_{sym} = \Sigma_{hkl} \Sigma_i | I_{hkl,i} - \langle I_{hkl} \rangle | / \Sigma_{hkl} \Sigma_i I_{hkl,i}$ and $R_{pim} = \Sigma_{hkl} (1/(n-1))^{1/2} \Sigma_i | I_{hkl,i} - \langle I_{hkl} \rangle | / \Sigma_{hkl} \Sigma_i I_{hkl,i}$, where $I_{hkl,i}$ is the scaled intensity of the $i^{th}$ measurement of reflection h, k, l, $\langle I_{hkl} \rangle$ is the average intensity for that reflection, and $n$ is the redundancy.
[d] $CC_{1/2}$ = Pearson correlation coefficient between two random half datasets.
[e] $R_{cryst} = \Sigma_{hkl} | F_o - F_c | / \Sigma_{hkl} | F_o |$ x 100, where $F_o$ and $F_c$ are the observed and calculated structure factors, respectively.
[f] $R_{free}$ was calculated as for $R_{cryst}$, but on a test set comprising 5-10% of the data excluded from refinement.
[g] Macromolecule atoms include protein and N-glycans.
[h] Ligands include nucleotides, PO4, glycerol, ethylene glycol, acetate, tartrate, and citrate.

**Table S2. Protein stability prediction of PLD3 mutants**

| | ddG (Kcal/mol) |
|---|---|
| mPLD3-L306P | -1.46 |
| mPLD3-I163M | -1.37 |
| mPLD3-V230M | -0.86 |

Protein stability was predicted by I-Mutant Suite [3].

ddG Value:

dG(mutant) − dG(WildType) in kcal/mole

Binary Classification:

ddG<0: Decrease Stability

ddG>0: Increase Stability

Ternary Classification:

ddG<−0.5: Large Decrease of Stability

ddG>0.5: Large Increase of Stability

−0.5<=ddG<=0.5: Neutral Stability
